# A Genome-wide CRISPR Screen Identifies WDFY3 as a Novel Regulator of Macrophage Efferocytosis

**DOI:** 10.1101/2022.01.21.477299

**Authors:** Jianting Shi, Xun Wu, Ziyi Wang, Fang Li, Yujiao Meng, Rebecca M. Moore, Jian Cui, Chenyi Xue, Katherine R. Croce, Arif Yurdagul, John G Doench, Wei Li, Konstantinos S. Zarbalis, Ira Tabas, Ai Yamamoto, Hanrui Zhang

## Abstract

Phagocytic clearance of dying cells, termed efferocytosis, must occur efficiently to maintain homeostasis and prevent disease. Yet, our understanding of this important biological process remains incomplete. To search for novel regulators of efferocytosis, we performed a FACS-based genome-wide CRISPR knockout screen in primary murine macrophages. We identified a novel role for WDFY3 in efferocytosis by macrophages. WDFY3 deficiency in macrophages specifically impaired uptake, not binding, of apoptotic cells due to defective actin depolymerization. We further revealed that WDFY3 directly interacts with GABARAP, thus facilitating LC3 lipidation and subsequent lysosomal acidification to permit the degradation of apoptotic cell components. Although the C-terminus of WDFY3 was sufficient to rescue impaired degradation, full-length WDFY3 is still required for regulating uptake. Finally, WDFY3 is required for efficient efferocytosis *in vivo* in mice and in primary human macrophages. The work expands our knowledge of the mechanisms of macrophage efferocytosis, and more broadly, provides a general strategy for genome-wide CRISPR screen to interrogate complex functional phenotypes in primary macrophages.

**Highlights:**

- Functional readout for pooled genome-wide CRISPR screen in primary macrophages.
- WDFY3 is discovered as a regulator of macrophage efferocytosis *in vitro* and *in vivo*.
- WDFY3 deficiency led to impaired uptake, as opposed to binding, of apoptotic cells due to defective actin depolymerization.
- WDFY3 directly interacts with GABARAP, facilitating LC3 lipidation and subsequent lysosomal acidification to permit the degradation of apoptotic cell components.
- C-terminal WDFY3 is sufficient to regulate the degradation of engulfed apoptotic cells while full-length WDFY is required for regulating uptake.

## Introduction

Phagocytic clearance of dead or dying cells by phagocytes, a process known as efferocytosis, is important in embryogenesis and development, and the resolution of pathological events^1–4^. Impaired efferocytosis lessens the effective clearance of dying cells, causing secondary necrotic cell death and damages^1–4^. Efferocytosis is performed by macrophages and to a lesser extent by other professional phagocytes (such as monocytes and dendritic cells), non-professional phagocytes and specialized phagocytes^1^. Because of the fundamental role of efferocytosis, dysregulation of this process is associated with many pathological states, including autoimmune diseases, atherosclerosis, and cancers^2^. Given the importance of this biological process and the therapeutic potential of targeting genes regulating efferocytosis, identifying novel regulators and mechanisms of this biological process has broad impacts on many diseases relevant to defective efferocytosis^5–8^.

Hypothesis-driven approaches have successfully identified many key regulators for the removal of dying cells via efferocytosis^1–4^. Yet, an unbiased approach to screening regulators of efferocytosis of apoptotic cells (ACs) on a genome-wide scale is lacking. Unbiased screenings allow the identification of new regulators from diverse and unexpected gene classes. Genetic screens of efferocytosis of ACs have been performed in *Drosophila*^9^, but not in mammalian cells. Genome-wide CRISPR knockout screens have recently been applied in macrophages differentiated from human U937 myeloid leukemia cell line to identify regulators for phagocytosis of diverse substrates, including zymosan, myelin, red blood cells with or without opsonization, and beads ranging from 0.3 μm to 4 μm^10, 11^; and in J774 murine macrophage-like cell line to identify regulators for phagocytosis of cancer cells^12^, illuminating both universal and substrate-specific principles of phagocytosis. However, a screening platform using ACs as the substrates and in primary macrophages is critical because efferocytosis involves AC-specific recognition receptors^13^, stiffness and size-dependent engulfment mechanisms^14^, and cellular response to degradation^4^, all of which cannot be recapitulated by phagocytosis of beads. In addition, immortalized or tumor-derived monocytic cell lines often lack physiological relevance to fully resemble the spectrum of physiological function in primary macrophages^15^.

To address this gap, we established and performed a pooled genome-wide CRISPR knockout screen for efferocytosis in primary murine bone marrow-derived macrophages (BMDM) derived from the Rosa26-Cas9 knock-in mice constitutively expressing Cas9 endonuclease. Our screen has successfully identified well-known key regulators responsible for the recognition and uptake of ACs, supporting the screen’s performance. Individual validation of the strongest hits has uncovered WDFY3 (WD repeat and FYVE domain containing 3), also known as Alfy (Autophagy-linked FYVE Protein), as a novel regulator previously not implicated in the regulation of efferocytosis or phagocytosis. We further uncovered the novel mechanisms by which WDFY3 regulates the uptake and degradation of ACs during efferocytosis and demonstrated the role of WDFY3-mediated efferocytosis in mice *in vivo* and in primary human macrophages *in vitro*. Our study also establishes a broadly-applicable platform for the genome-wide screen of complex functional phenotypes in primary macrophages for unbiased novel discoveries.

## Results

### A pooled, FACS-based genome-wide CRISPR knockout screen in primary macrophages identified known and novel regulators of macrophage efferocytosis

Genome-wide forward genetic screens have the capacity to examine a biological process in an unbiased manner and allow for novel discoveries. We first determined the proper cell types for a genome-wide CRISPR screen of macrophage efferocytosis. Human monocytic cell lines, including U937 and THP-1, can be differentiated to macrophage-like cells, which have previously been used for genome-wide screening^10, 11^. Yet, we confirmed that U937 and THP-1 derived macrophages were not a proper model for screening of efferocytosis as monocytic cell line-derived macrophages showed poor efferocytosis capacity. Specifically, upon up to 24 hours of AC incubation, only less than 1-3 % of either U937-derived or THP-1-derived macrophages were able to engulf ACs (**Supplementary Fig. 1a**). The results highlight the importance of using physiologically relevant primary macrophages for screening of efferocytosis regulators.

We thus leveraged the Rosa26-Cas9 knock-in mice constitutively expressing Cas9 endonuclease (JAX #026179)^16^ and established a workflow for CRISPR gene editing in primary bone-marrow-derived macrophages (BMDMs). Specifically, lentiviral gRNA libraries were transduced to isolated bone marrow (BM) cells, which were then differentiated to BMDMs using L cell-conditioned media that provide macrophage colony-stimulating factor (M-CSF) for macrophage differentiation. As illustrated in **Fig. 1a** and **Fig. 1b**, for each replicate, 400-500 million BM cells were isolated and seeded. The lentiviral Brie library^17^ (Addgene 73633) including 78,637 gRNAs targeting 19,674 mouse genes and 1,000 non-targeting control gRNAs was transduced on day 1 with a low MOI to ensure that majority of the BM cells integrate one viral particle for gene editing of a single gene. 48 hours after transduction, puromycin was applied to select BM cells with successful lentiviral integration.

**Fig. 1.**
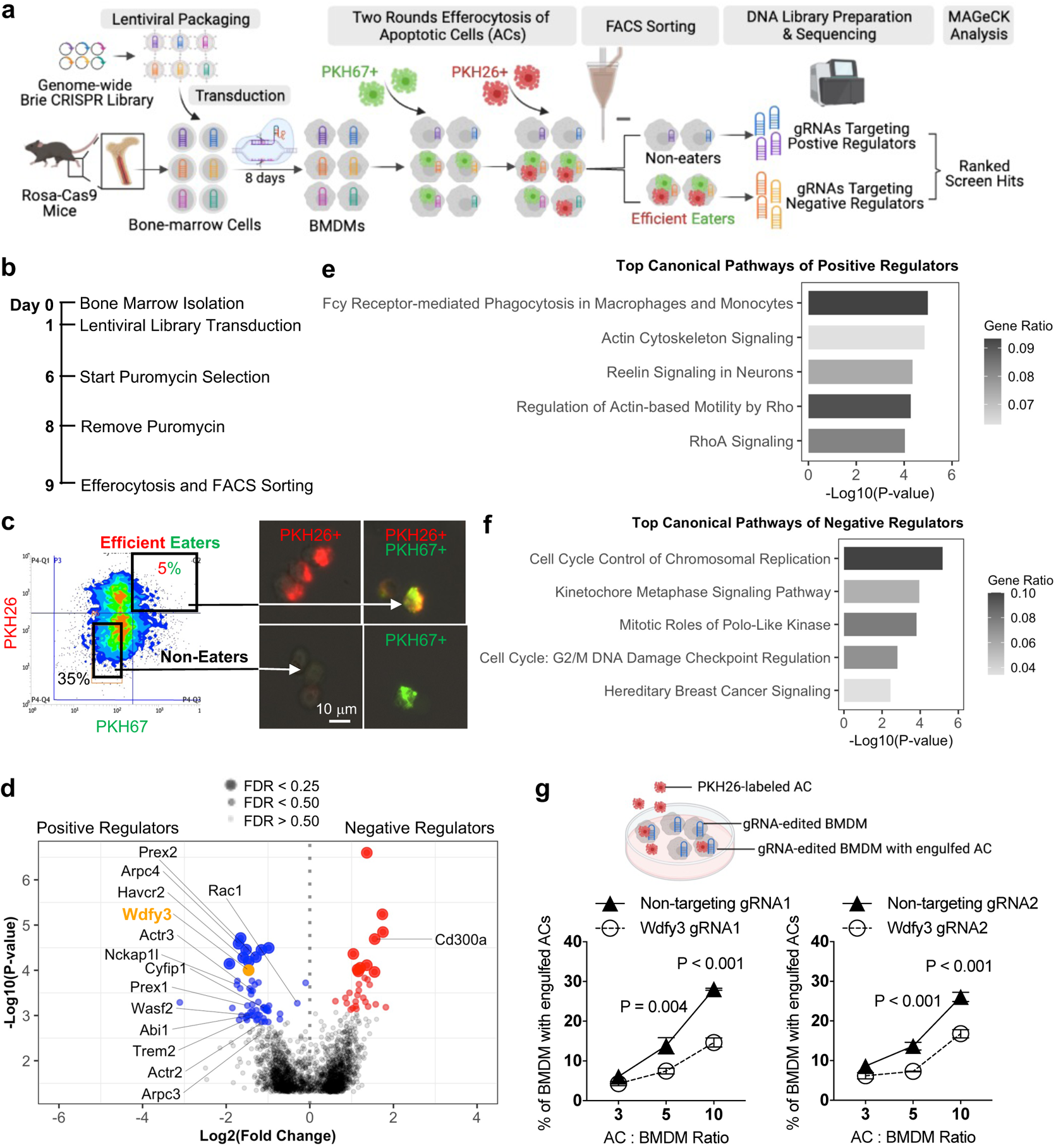
A pooled, FACS-based genome-wide CRISPR knockout screen in primary macrophages identified known and novel regulators of macrophage efferocytosis. **(a)** Schematics of the CRISPR screen workflow. **(b)** Timeline of bone marrow isolation, lentiviral library transduction, puromycin selection, efferocytosis, and cell sorting. **(c)** Visualization of gating strategy for separation of non-eaters and efficient eaters. Successful separation was confirmed by fluorescent microscopy. **(d)** Volcano plot highlights the top-ranked screen hits that are known positive and negative regulators of macrophage efferocytosis. **(e)** Canonical pathways enriched in top-ranked positive regulators by Ingenuity Pathway Analysis (IPA). **(f)** Canonical pathways enriched in top-ranked negative regulators by IPA. **(g)** Validation of *Wdfy3* as a positive regulator required for macrophage efferocytosis (n = 4 independent experiments).

The success of the screening relies on the effective enrichment of macrophages with high vs. low efferocytosis capacity. Since efferocytosis is a binary event, to facilitate an effective separation and enrichment, we performed two rounds of efferocytosis sequentially. Specifically, human Jurkat cells (~10 μm in diameter), an acute T cell leukemia cell line routinely used for *in vitro* efferocytosis assays, were treated with staurosporine to induce apoptosis, followed by labeling with fluorescent linkers, PKH67 (Ex/Em: 490/502 nm) or PKH26 (Ex/Em: 551/567 nm), that incorporate into the cell membrane lipid bilayer for stable labeling of the cell membrane. BMDMs were first incubated with PKH67-labeled ACs at a ratio of 5:1 for AC : BMDM and allowed for efferocytosis. After 45 minutes, the unbound PKH67-labeled ACs were washed away and BMDMs were cultured for two hours without ACs to allow degradation of the engulfed cargo. BMDMs were then fed with PKH26-labeled ACs also at a ratio of 5:1. After 90 minutes, unbound ACs were washed away and BMDMs were collected for flow cytometry sorting to separate the BMDMs that engulfed both PKH67^+^ and PKH26^+^ ACs, i.e., the efficient eater (~5%), and BMDMs that did not engulf any ACs, i.e., the non-eater (**Fig. 1c and Supplementary Fig. 1b**). Two independent replicates were performed (**Supplementary Fig. 1c**). For each replicate, efferocytosis was performed in ~80 million BMDMs on day 9 (**Supplementary Fig. 1b**). After sorting, we obtained ~3 million efficient eaters and ~16 million non-eaters. We have also collected 40 million BMDMs on day 9 without performing efferocytosis, i.e. the input samples (**Supplementary Fig. 1b**).

We sequenced the sorted non-eaters, efficient eaters, and the input samples for each of the two replicates and performed MAGeCK analysis^18–21^ to identify the top hits. We analyzed three comparisons: input vs. non-eaters (**Supplementary Table 1**), input vs. efficient eaters (**Supplementary Table 2**), non-eaters vs. efficient eaters (**Supplementary Table 3**). We expect that the comparison of input vs. non-eaters will identify positive regulators whose knockout impairs efferocytosis, while the comparison of input vs. efficient eaters will identify negative regulators whose knockout enhances efferocytosis. The comparison of non-eaters vs. efficient eaters likely further improves the power to identify enriched gRNAs. As expected, the analysis comparing non-eaters vs. efficient eaters was able to identify more known regulators (**Fig. 1d, Supplementary Table 3** for the complete MAGeCK output). Non-targeting gRNAs did not show enrichment in either sample (**Supplementary Fig. 1d**).

The non-eaters are expected to enrich for gRNAs targeting positive regulators essential for efferocytosis, i.e., knockout would impair efferocytosis. Indeed, we identified many genes involved in actin polymerization that is known to be essential for phagocytic cup formation, including *Rac1*, four members of the five-subunit SCAR/WAVE complex (*Nckap1l*, *Wasf2*, *Abi1*, *Cyfip1*) and five members of the seven-subunit ARP2/3 complex (*Actr2*, *Actr3*, *Arpc3*, and *Arpc4*) (**Fig. 1d**). We performed pathway analysis using Ingenuity Pathway Analysis (IPA). The top-ranked positive regulators (negative score < 0.002, 163 genes) were enriched for pathways including “Fcγ Receptor-mediated Phagocytosis in Macrophages and Monocytes”, “Actin Cytoskeleton Signaling” etc. (**Fig. 1e** and **Supplementary Table 4**), supporting the screening performance in identifying well-known positive regulators. The results also showed that many, but not all, genes involved in actin cytoskeleton remodeling and general phagocytosis are among the most highly ranked screen hits (**Supplementary Fig. 2**).

Using high-content imaging analysis, we selectively validated *Arpc4* (top-2 ranked) and *Nckap1l* (top-14 ranked) using the gRNAs from the original screening library. gRNAs targeting *Arpc4* or *Nckap1l* led to ~50% reduction in the efferocytosis of PKH26-labeled ACs by BMDM (**Supplementary Fig. 1e).** *Hacvr2*, also known as TIM3, is one of the PtdSer-specific receptors involved in AC recognition and efferocytosis^22^. *Hacvr2* was ranked at top-7 and was also validated with ~30% reduction in efferocytosis capacity (**Supplementary Fig. 1e**).

The efficient eaters are expected to enrich for gRNAs targeting negative regulators, i.e. knockout would enhance efferocytosis. Efferocytosis needs to be tightly controlled and there are very few known negative regulators. While this manuscript is being prepared, the top-2 ranked hit for negative regulators, *Cd300a* (**Fig. 1d**), was identified as a novel negative regulator^23^. Specifically, the binding of an AC with Cd300a and the activation of downstream signaling suppresses efferocytosis by myeloid cells, thus the blockage of *Cd300a* enhanced efferocytosis^23^. We were also able to validate the results in BMDM using a gRNA targeting *Cd300a* (**Supplementary Fig. 1e**). Pathway analysis of top-ranked negative regulators (positive score <0.001 for a total of 96 genes) implies that genes involved in cell cycle control and chromosomal replication were enriched for top hits for negative regulators (**Fig. 1f** and **Supplementary Table 5**).

The screen has revealed many top-ranked hits that promise to inform novel biology and warrant further validation and functional interrogation. Among the top hits for positive regulators, *Wdfy3*, the top-10 ranked, is a novel one that has not been implicated in the regulation of efferocytosis or phagocytosis, nor identified in previous screens in non-mammalian cells or using other substrates (**Supplementary Fig. 2 and Supplementary Table 6**). Using two individual gRNAs, one from the Brie library and one designed independently, and quantitative imaging analysis, we validated that knockout of *Wdfy3* in BMDMs led to impaired efferocytosis of PKH26-labeled ACs (**Fig. 1g**). The defects were more significant when BMDMs were challenged with higher AC to BMDM ratio, i.e., a condition mimicking high-burden efferocytosis.

Altogether, our group is the first to establish a CRISPR screen for regulators of efferocytosis, a complex functional phenotype, in primary macrophages at genome-wide coverage. WDFY3 is uncovered as a novel regulator that we focus on determining the mechanisms.

### WDFY3 deficiency led to impaired uptake, as opposed to binding, of apoptotic cells due to defective actin depolymerization

*WDFY3* encodes a highly conserved, large 400 kDa protein with 3526 amino acids. Similar to mouse^24^, *WDFY3* mRNA is the most abundantly expressed in the brain (**Supplementary Fig. 3a**). Consistently, at the single-cell level, *WDFY3* is most abundantly expressed in multiple brain cell types (**Supplementary Fig. 3b**). Among immune cells, *WDFY3* is abundantly expressed in myeloid cells, including macrophages, neutrophils, and monocytes, but not T cells (**Supplementary Fig. 3b** and **Supplementary Fig. 3c**).

To further validate the role of *Wdfy3* knockout in efferocytosis *ex vivo*, we obtained *Wdfy3^fl/fl^* mice created by insertion of two *loxP* sites flanking exon 5 on a 129/SvEv x C57BL/6 background as previously described^24^. Myeloid-specific *Wdfy3* null mice were generated by breeding *Wdfy3^fl/fl^* mice with LysMCre mice (JAX 004781), i.e. *LysMCre^+/−^Wdfy3^fl/fl^* mice (Cre^+^) while using *LysMCre^−/−^Wdfy3^fl/fl^* littermates (Cre^−^) as the controls (as illustrated in **Fig. 2a)**. We confirmed efficient knockout by western blotting of WDFY3 in BMDMs from the Cre^+^ mice (**Fig. 2b**). Although global deletion of *Wdfy3* led to perinatal lethality^24^, myeloid-specific loss of *Wdfy3* did not affect body weight (**Supplementary Fig. 4a**) or organ weight, including that of heart, liver, and spleen (**Supplementary Fig. 4b)**. Moreover, the mice did not show changes in circulating levels of neutrophils and monocytes, confirming that myelopoiesis was not affected (**Supplementary Fig. 4c**).

**Fig. 2.**
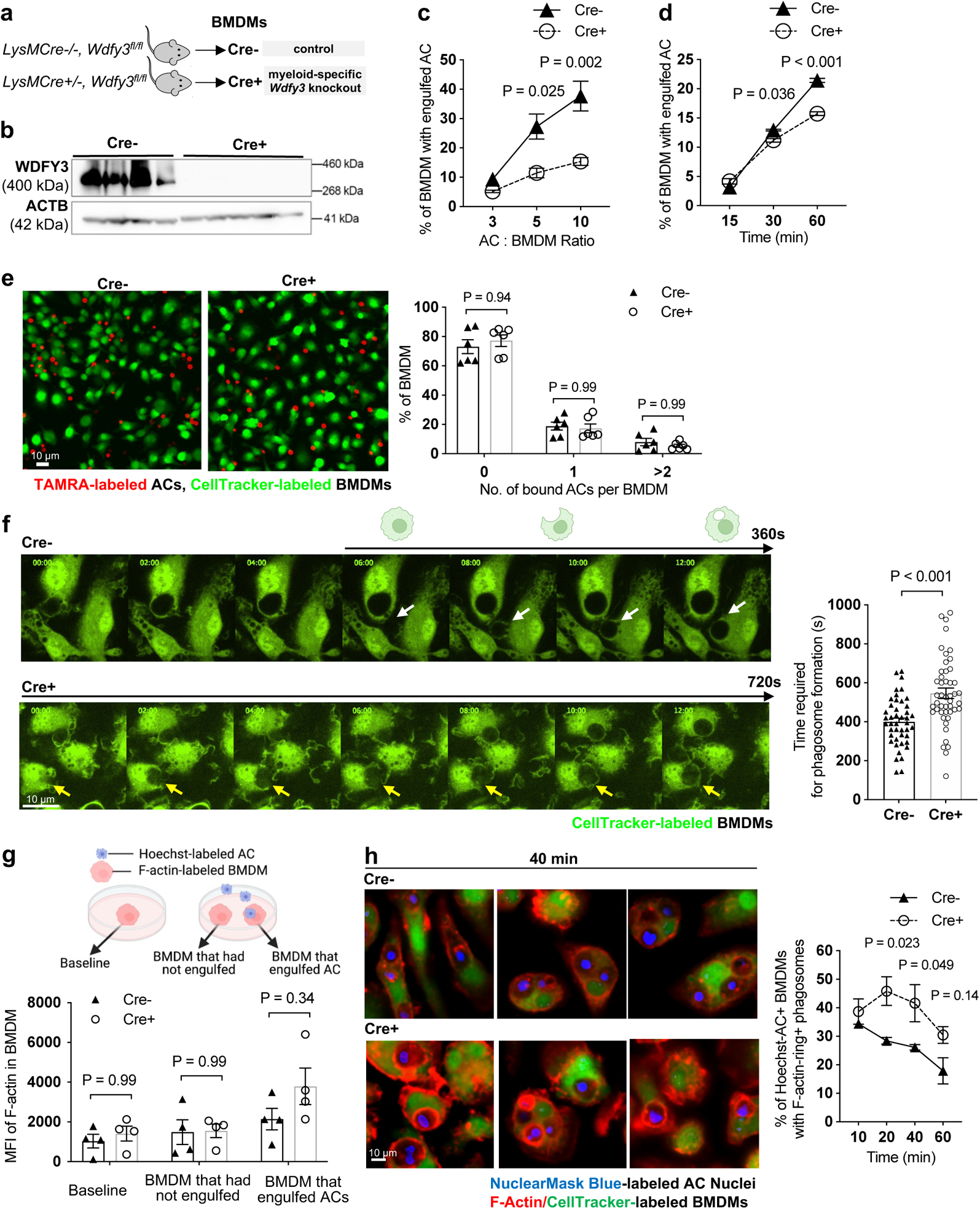
WDFY3 deficiency led to impaired uptake, as opposed to binding, of apoptotic cells (ACs) due to defective actin depolymerization. **(a)** Schematics of breeding LysMCre mice with *Wdfy3^fl/fl^* mice to obtain myeloid-specific knockout of *Wdfy3*. **(b)** Validation of efficient knockout in BMDMs by Western Blot of WDFY3 (n = 4 biological replicates; the plot shown is a representative image of three independent experiments). **(c)** Cre^−^ and Cre^+^ BMDMs were incubated with PKH26-labeled ACs at various AC : BMDM ratios of 3:1, 5:1, 10:1 respectively for 1 hour and analyzed by flow cytometry (n = 3 biological replicates, each from the average of 2 technical replicates). **(d)** Cre^−^ and Cre^+^ BMDMs were incubated with PKH26-labeled ACs at various time points of 15 min, 30 min, and 60 min at a AC : BMDM ratio of 5:1 and analyzed by flow cytometry (n = 3 technical replicates). **(e)** Cre^−^ and Cre^+^ BMDMs were pre-treated with cytochalasin D for 30 min to block polymerization and elongation of actin, thus testing the binding of ACs with BMDMs. The treated BMDMs were then incubated with TAMRA-stained apoptotic mouse thymocytes at 37 °C for 30 min and then extensively washed with PBS to remove unbound ACs for imaging and quantification after fixation (n = 6 biological replicates). **(f)** Cre^−^ and Cre^+^ BMDMs were stained with CellTracker and incubated with ACs. Efferocytosis of ACs by BMDMs were observed using time-lapse confocal microscopy. The time required for phagosome formation was measured and recorded (n = 4 biological replicates, each data point represents one BMDM with engulfed ACs). **(g)** F-actin labeled by siR-actin in Cre^−^ and Cre^+^ BMDMs was quantified by flow cytometry (n = 4 biological replicates, each from the average of 3 technical replicates). **(h)** BMDMs were stained with CellTracker and siR-actin, then incubated with NuclearMask Blue-labeled apoptotic Jurkat cells for various time points (10 min, 20 min, 40 min, and 60 min). For each time point, unbound ACs were removed and BMDMs were fixed. BMDMs were imaged and the percentage of BMDMs with engulfed cargos surrounded by F-actin rings in all BMDMs with engulfed cargos was quantified (n = 3 biological replicates, data are representative of two independent experiments).

We used flow cytometry to quantify the percentage of BMDMs with engulfed PKH26-labeled ACs. With lower AC : BMDM ratio or at relatively early time points, efferocytosis of Cre^−^ and Cre^+^ BMDMs appeared similar (**Fig. 2c** and **Fig. 2d**). The defects were more significant with a high ratio of AC : BMDM that resembles high-burden efferocytosis (**Fig. 2c**). Consistently, with an AC : BMDM ratio at 5:1, the defective efferocytosis in Cre^+^ BMDMs was the most significant at later time points (**Fig. 2d**), also supporting more pronounced defects over prolonged periods of challenges.

Efferocytosis involves the finding, recognition and binding, uptake, and subsequently the degradation of the engulfed cargos^1, 2^. Our screen is designed to identify regulators essential for the binding and/or uptake of ACs. The screen will not identify genes only regulating the degradation without affecting the binding or uptake of ACs. The screen will also not identify genes specifically responsible for the chemotactic cues termed “find-me” signals because the pooled design masks the defective secretion by a small subset of edited cells. Yet, genes regulating the finding or degradation of ACs can be identified if they also regulate the binding or uptake of ACs. We next set out to determine the molecular steps regulated by WDFY3. We first aimed at determining if *Wdfy3* knockout affected binding and/or uptake during efferocytosis. TAMRA-labeled apoptotic murine thymocytes were incubated with CellTracker-labeled BMDMs pretreated with cytochalasin D that prevents actin polymerization thus the uptake of ACs. Following incubation, unbound ACs were washed away and BMDMs were fixed and imaged. The numbers of TAMRA-labeled ACs bound with each BMDM were counted and the percentage of BMDMs with none, one, or two and more bound ACs was quantified for Cre^−^ and Cre^+^ BMDMs. The results support that *Wdfy3* knockout did not affect the ability of BMDMs to bind ACs (**Fig. 2e**), suggesting that the uptake, as opposed to binding, of ACs was impaired due to *Wdfy3* deficiency.

Indeed, time-lapse live-cell imaging confirmed that the time required for complete internalization of ACs was longer in *Wdfy3* knockout BMDM compared with control (**Fig. 2f**), suggesting delayed phagosome formation. Phagosome formation during phagocytosis of large particles requires the coordination of actin polymerization and depolymerization, permitting the continual restructuring of the actin cytoskeleton^14^. It has been described that complete internalization of the cargo is synchronized with actin depolymerization, allowing subsequent phagosome maturation^25^ (as also visualized in **Supplementary Video 1**). We thus asked if *Wdfy3* knockout affected actin polymerization and/or depolymerization. We labeled BMDMs with siR-actin, a fluorogenic, cell-permeable probe based on an F-actin binding natural product jasplakinolide, and determined F-actin levels at baseline and upon efferocytosis of Hoechst-labeled ACs by flow cytometry. F-actin signals at baseline were similar between Cre^−^ and Cre^+^ BMDMs (**Fig. 2g**, **left panel**). Upon efferocytosis, BMDMs that had not engulfed ACs also showed comparable F-actin levels between Cre^−^ and Cre^+^ BMDMs (**Fig. 2g**, **middle panel**). Yet, in BMDMs that engulfed ACs, *Wdfy3* knockout BMDMs showed higher F-actin signals (**Fig. 2g**, **right panel**, P = 0.34). Although not statistically significant, the observed trend led us to hypothesize that potential defects in actin depolymerization exist in *Wdfy3* knockout BMDMs. Indeed, in *Wdfy3*-deficient BMDMs that had successfully internalized an AC, we observed that many engulfed ACs were surrounded by F-actin rings (**Fig. 2h**). Confirming our subjective observations, the percentage of BMDMs with F-actin surrounded cargos over all BMDMs that had engulfed ACs was greater in Cre^+^ vs. Cre^−^ BMDMs (**Fig. 2h**). In Cre^−^ control BMDMs, the percentage of BMDMs with F-actin surrounded cargos was the highest at 10 min after adding ACs and then decreased over time (**Fig. 2h**). Yet, in Cre^+^ BMDMs, the percentage further increased and peaked at 20 min after adding ACs and remain greater than the percentage in Cre^−^ BMDMs (**Fig. 2h**), supporting defective actin depolymerization.

Thus, defective actin depolymerization in *Wdfy3* knockout macrophages led to impaired uptake and delayed phagosome formation during efferocytosis. The defects were specific to efferocytosis of ACs because the phagocytosis of other substrates, including beads of different sizes (4 μm and 10 μm, **Supplementary Fig. 5a** and **Supplementary Fig. 5b**), sheep red blood cells (RBCs) that were untreated, stressed by heat treatment, or IgG-opsonized (**Supplementary Fig. 5c**), zymosan particles (500 nm, **Supplementary Fig. 5d**), was not impaired in *Wdfy3* knockout BMDM. Consistently, previous screens using the above-mentioned substrates in U937 monocytic line-derived macrophages^11^, or using cancer cells in J774 macrophages^12^ did not uncover *Wdfy3* as a hit (**Supplementary Fig. 2** and **Supplementary Table 6**). Thus, we discovered and validated a novel regulator specifically required for the uptake of ACs during efferocytosis.

We have also validated the role of *Wdfy3* in macrophage efferocytosis in *Wdfy3^fl/fl^* mice generated by the Knock-Out Mouse Project (KOMP) with two *loxp* sites flanking exon 8, and maintained on C57BL/6N background^26^. Breeding to LysMCre mice led to efficient knockout of WDFY3 though a small amount of residual protein remained detectable (**Supplementary Fig. 6a**). Consistently, we have observed impaired uptake of ACs in Cre^+^ BMDMs (**Supplementary Fig. 6b**), further confirming that the role of *Wdfy3* in macrophage efferocytosis is independent of the genetic strain or specific gene-inactivating mutation of the mouse models.

### WDFY3 deficiency led to impaired degradation of engulfed ACs

Sustained accumulation of periphagosomal F-actin prevents efficient phagosome-lysosome fusion^25^. We thus reasoned that defective actin depolymerization may impair the degradation of the engulfed cargos. To test the hypothesis, we determined the degradation of the engulfed ACs by Cre^+^ and Cre^−^ BMDMs. We first incubated BMDMs with PKH26-labeled ACs for efferocytosis. After 60 minutes of incubation, unbound ACs were washed away and BMDMs were returned to the incubator for three hours to allow degradation of the engulfed cargos. BMDMs were then fixed and imaged. We counted the percentage of AC^+^ BMDMs that showed non-fragmented PKH26 staining implicating impaired degradation. Indeed, the percentage of BMDMs with non-fragmented ACs was greater in Cre^+^ vs. Cre^−^ BMDMs (**Fig. 3a**), confirming impaired degradation in *Wdfy3*-deficient BMDMs.

**Fig. 3.**
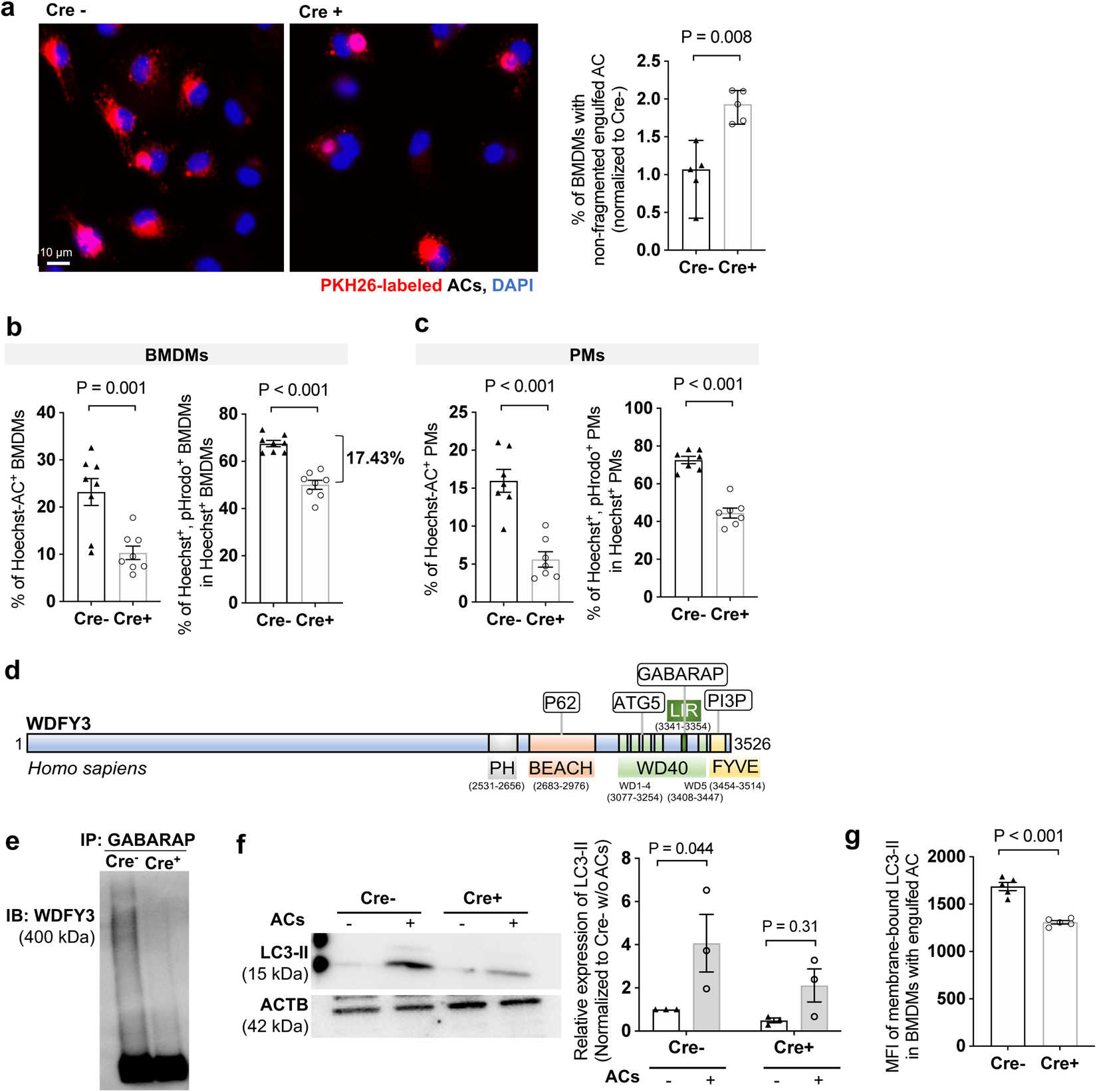
WDFY3 deficiency led to defects in LC3-associated phagocytosis (LAP) and the degradation of engulfed ACs. **(a)** Cre^−^ and Cre^+^ BMDMs were incubated with PKH26-labeled ACs for one hour. After washing away the unengulfed ACs, BMDMs were placed back to the incubator for another three hours. BMDMs were then fixed and imaged. The percentage of BMDMs showing non-fragmented PKH26 signals in the total number of PKH26^+^ BMDMs was quantified (n = 5 biological replicates, each from the average of 3 technical replicates). Cre^−^ and Cre^+^ BMDMs **(b)** and PMs **(c)** were incubated with ACs labeled by Hoechst, which stains DNA and is pH-insensitive, and pHrodo, which is pH-sensitive and shows fluorescent signal only under an acidified environment in the phagolysosome. The percentage of Hoechst^+^ BMDMs indicates uptake. The percentage of Hoechst^+^/pHrodo^+^ BMDMs in Hoechst^+^ BMDMs indicates acidification of the engulfed cargos (n = 8 biological replicates, each from the average of 2 technical replicates). **(d)** Schematics of known functional domains and binding partners of human WDFY3. **(e)** The interaction between WDFY3 and GABARAP was assessed by co-immunoprecipitation. Cre^−^ and Cre^+^ BMDMs cell lysate were incubated with anti-GABARAP antibody-conjugated agarose beads. Beads-bound proteins were detected with anti-WDFY3 antibodies. **(f)** Cre^−^ and Cre^+^ BMDMs were incubated with ACs for one hour. Unbound ACs were washed away and BMDMs were collected for measurement of LC3-II by western blot (n = 3 independent experiments). **(g)** BMDMs were incubated with Hoechst-labeled ACs to allow efferocytosis. After removal of unbound ACs, BMDMs were collected, fixed, and treated with digitonin to remove non-membrane bound LC3, and then immunostained for LC3 that is lipidated and membrane-bound. LC3-II staining was then quantified by flow cytometry for BMDMs that had engulfed Hoechst-labeled ACs (n = 5 biological replicates).

To dissect if the impaired degradation in *Wdfy3* knockout BMDMs was also linked with impaired lysosomal acidification, we dual-labeled ACs with Hoechst that stains DNA and is pH-insensitive, and pHrodo-Red that is pH-sensitive and shows fluorescent signals only under an acidified environment in the phagolysosome. *Wdfy3* knockout BMDMs showed impaired efferocytosis of Hoechst-labeled ACs (**Fig. 3b, left panel**), consistent with the results when using PKH26-labeled ACs (**Fig. 2c**). For BMDMs with engulfed Hoechst^+^ ACs, the percentage of pHrodo^+^/Hoechst^+^ BMDMs in Hoechst^+^ BMDMs is lower in Cre^+^ BMDMs vs. Cre^−^ BMDMs, supporting impaired acidification in *Wdfy3* knockout BMDMs (**Fig. 3b, right panel**). We observed consistent results using peritoneal macrophages (PMs) (**Fig. 3c**). Thus, WDFY3 is required for both the uptake and the degradation of engulfed ACs during efferocytosis, and *Wdfy3*-deficiency led to impaired acidification of the phagolysosome.

### WDFY3 deficiency led to defects in LAP

We next asked if the impaired degradation in *Wdfy3* knockout BMDMs was merely a consequence of the defects in actin depolymerization during phagosome formation or mediated by other potentially independent mechanisms. We first considered whether WDFY3 is involved in LC3-associated phagocytosis (LAP), a process by which LC3-II conjugation to phagosomes enables phagosome-lysosome fusion and AC corpse degradation^27–32^. Our hypothesis is built on the known role of WDFY3 in autophagic clearance of aggregated proteins, i.e. aggrephagy^33, 34^. Specifically, The C-terminus of both mouse and human WDFY3 contains several functional domains (as illustrated in **Fig. 3d**)^35^. Co-immunoprecipitation and colocalization studies indicated that WDFY3 scaffolds a complex containing the p62-positive, ubiquitinated, aggregation-prone protein and the core autophagy proteins ATG5, ATG12, ATG16L1 and LC3/GABARAP^33, 36^. The human ortholog of the yeast *Atg8* includes the LC3 family (LC3A, LC3B, LC3B2 and LC3C) and the GABARAP family (GABARAP, GABARAPL1 and GABARAPL2).

During aggrephagy, the WD40 repeats of WDFY3 are essential for its colocalization and interaction with ATG5^33^. The ATG5-ATG12 complex is required for an early stage of autophagosome formation, and together with the membrane-bound ATG16L1 facilitate the conjugation of LC3/GABARAP proteins to phosphatidylethanolamine, i.e. LC3 lipidation to form LC3-II, for autophagosome formation^37, 38^. Recent work using HEK293T cells further revealed that WDFY3 has a conserved LIR (LC3-interacting region) motif in its WD40 region that directly binds to GABARAP, responsible for its recruitment to LC3B during aggrephagy^36^.

We first set out to determine if endogenous WDFY3 interacts with GABARAP in macrophages. Whole-cell lysates from Cre^−^ and Cre^+^ BMDMs were incubated with anti-GABARAP antibodies and were immunoprecipitated using protein A/G beads. WDFY3 can be found in a complex with endogenous GABARAP in Cre^−^ BMDMs, confirming WDFY3 and GABARAP interactions (**Fig. 3e**). No precipitation was observed in *Wdfy3* knockout BMDMs, confirming the specificity of the antibody (**Fig. 3e**).

We thus reasoned that WDFY3 interacts with GABARAP, regulating the recruitment and lipidation of LC3 during LAP for subsequent cargo degradation. Consistent with previous literature^39^, AC engulfment led to increased LC3-II as determined by western blot (**Fig. 3f**). The increase was blunted in *Wdfy3*-deficient BMDMs (**Fig. 3f**). We further confirmed the results using a flow cytometry-based assay as described^40^. Specifically, BMDMs were incubated with Hoechst-labeled ACs to allow efferocytosis. After removal of unbound ACs, BMDMs were collected, fixed, and treated with digitonin to remove non-membrane bound LC3, and then immunostained for LC3 that is lipidated and membrane-bound, i.e. LC3-II. As quantified by flow cytometry, for BMDMs that had engulfed Hoechst-labeled ACs, *Wdfy3*-deficient BMDMs had lower membrane-bound LC3-II (**Fig. 3g**), supporting impaired LC3 lipidation.

Taken together, WDFY3 regulates LAP-mediated degradation of engulfed ACs through interacting with GABARAP and facilitating LC3 lipidation and the subsequent phagolysosomal degradation.

### A C-terminus fragment of WDFY3 is sufficient for regulating degradation yet the full-length protein is required for the AC uptake during efferocytosis

It has previously been shown that a 1000 amino acid C-terminus fragment, that contains the PH-BEACH, WD40, LIR, and FYVE domains of WDFY3 or the D. melanogaster ortholog, Bluecheese, was sufficient to enhance the degradation of aggregated proteins in otherwise wild-type cells^33, 41^. We therefore asked if this fragment was sufficient to regulate uptake and/or degradation during efferocytosis. We used lentiviral transduction to express C-terminal WDFY3 in both Cre^−^ and Cre^+^ BM cells that were then differentiated to BMDMs (as illustrated in **Fig. 4a**). Although expression of C-terminal WDFY3_(2543-3526)_ did not rescue the defective uptake in Cre^+^ BMDMs (**Fig. 4b**), it was sufficient to partly rescue the defects in the acidification of the engulfed ACs (**Fig. 4c**). Mechanistically, expression of C-terminal WDFY3 restored LC3 lipidation as quantified by both western blot (**Fig. 4d**) and flow cytometry (**Fig. 4e**). Thus, C-terminal WDFY3 is sufficient to facilitate LC3 lipidation and subsequent phagosome-lysosome fusion and phagolysosomal acidification, however, full-length WDFY3 is required for the regulation of efficient uptake during efferocytosis.

**Fig. 4.**
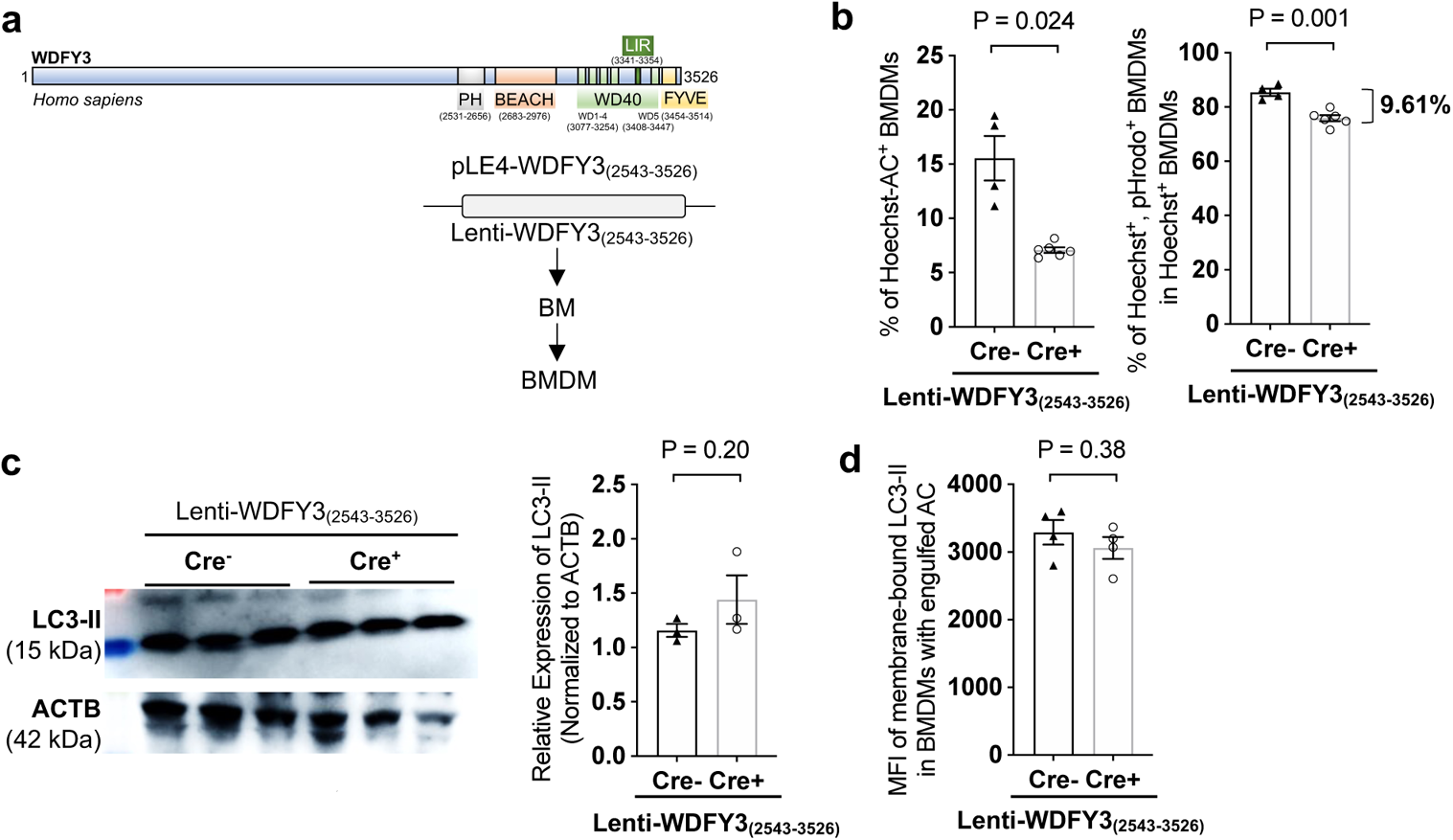
The C-terminal WDFY3 is sufficient for regulating degradation yet the full-length WDFY3 is required for the uptake of ACs during efferocytosis. **(a)** Schematics of lentiviral overexpression of C-terminal WDFY3 in BMDMs of Cre^−^ and Cre^+^ mice. **(b)** C-terminal WDFY3 did not restore uptake, yet partially rescued the defects in cargo acidification (n = 4 biological replicates, each from the average of 2 technical replicates). **(c)** and **(d)** C-terminal WDFY3 restored LC3-II levels as determined by western blot (n = 3 biological replicates, each from the average of 2 technical replicates) and by flow cytometry (n = 4 biological replicates, each from the average of 2 technical replicates).

### WDFY3 deficiency subtly affects the transcriptome of BMDMs without affecting macrophage differentiation

To gain an unbiased view of how *Wdfy3* knockout affects the transcriptomic signature of macrophages, we performed RNA-seq in Cre^−^ and Cre^+^ BMDMs (n = 4 male mice, **Supplementary Fig. 7**). We first confirmed that many receptors responsible for efferocytosis and phagocytosis, including *Fcgr1*, *Fcgr2b*, *Fcgr3, Mertk*, *Timd4*, and many macrophage marker genes, were expressed at similar levels between Cre^−^ and Cre^+^ BMDMs (**Supplementary Table 7**). Using a FDR-adjusted P value < 0.05 and absolute fold-change > 1.5, only a small number of genes were identified as differentially expressed (DE) between Cre^−^ and Cre^+^ BMDMs, i.e., 23 genes were upregulated while 31 genes were downregulated in Cre^+^ vs. Cre^−^ BMDMs (**Supplementary Fig. 7a** and **Supplementary Table 7**).

We reasoned that modest changes in the expression of genes belonging to the same pathway may imply functional impact. We thus performed gene-set enrichment analysis to determine which gene sets or pathways were enriched in upregulated or downregulated genes due to *Wdfy3* knockout. The upregulated genes in Cre^+^ BMDM were enriched for Human Reactome Pathways, including “IL-4 and IL-13 Signaling” and “Collagen Formation”, and GO Biological Process term “Regulation of Chemotaxis” (**Supplementary Fig. 7b** for representative plots, and **Supplementary Table 8 and Supplementary Table 9** for the complete GSEA output). The downregulated genes in Cre^+^ BMDM were enriched for Human Reactome Pathway “Peroxisomal Lipid Metabolism” and Gene Ontology (GO) Biological Process “Fatty Acid Catabolic Process” (**Supplementary Fig. 7c** for representative plots, and **Supplementary Table 10 and Supplementary Table 11** for the complete GSEA output). Overall, no clear pro-inflammatory or anti-inflammatory gene signatures were identified in Cre^+^ BMDMs.

We thus confirmed that: (1) Despite the profound role of WDFY3 in AC uptake and degradation, the observed transcriptomic modifications by *Wdfy3* knockout were only modest; (2) *Wdfy3* knockout did not affect macrophage maturation, as macrophage marker genes were not differentially expressed. We further confirmed that the percentage of F4/80^+^ macrophages in BMDMs and PMs was comparable between Cre^−^ and Cre^+^ mice (**Supplementary Fig. 8a**). Population doubling during BMDM differentiation was not different between Cre^−^ and Cre^+^ mice, supporting comparable differentiation and proliferation capacity (**Supplementary Fig. 8b**).

### Mice with myeloid *Wdfy3* deficiency show impaired efferocytosis *in vivo*

To determine if *Wdfy3* regulates efferocytosis *in vivo*, we performed two *in vivo* efferocytosis assays in Cre^−^ and Cre^+^ mice as illustrated in **Fig. 5a** (thymus efferocytosis) and **Fig. 5g** (PM efferocytosis).

**Fig. 5.**
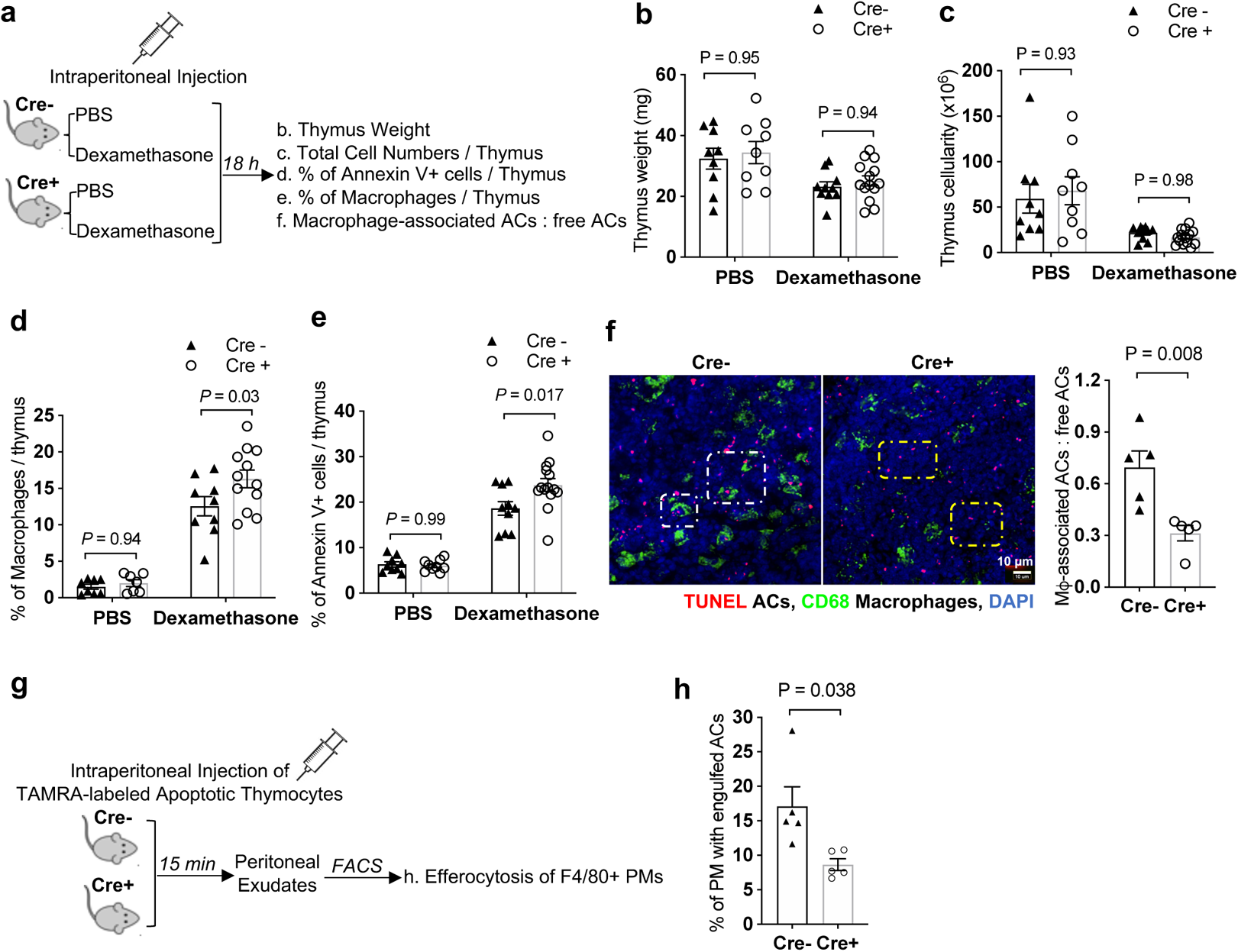
Mice with myeloid *Wdfy3* deficiency show impaired efferocytosis *in vivo*. **(a)** Schematics of experimental design for *in vivo* thymus efferocytosis assay. **(b)** Thymus weight. **(c)** Total number of cells per thymus. **(d)** Percentage of F4/80^+^ macrophages in the thymus determined by flow cytometry. **(e)** Percentage of Annexin V^+^ apoptotic cells per thymus determined by flow cytometry. A higher percentage implies impaired efferocytic clearance (n = 9 or 14 biological replicates for panel **b**, **c**, **d**, and **e**). **(f)** Thymic sections were stained with TUNEL for ACs, and CD68 for macrophages. The ratio of macrophage-associated TUNEL^+^ cells vs. free TUNEL^+^ cells was quantified and summarized. The white and yellow squares highlight the macrophage-associated and free TUNEL^+^ cells, respectively (n = 5 biological replicates). **(g)** Schematics of experimental design for *in vivo* peritoneal macrophage efferocytosis assay. **(h)** Peritoneal exudates were stained for F4/80 and the percentage of TAMRA^+^ peritoneal macrophages was determined by flow cytometry (n = 5 biological replicates).

For thymus efferocytosis (**Fig. 5a**), we treated Cre^−^ and Cre^+^ mice with dexamethasone that induces apoptosis of thymocytes, using PBS as the control. 18 hours after injection, thymi were isolated and weights were measured. The total number of cells per thymus was determined by dissociating one lobe of the thymus to count the cell number and then normalized to the weight of both lobes of the thymus. The dissociated cells were stained for Annexin V, a marker of apoptosis, and macrophage marker F4/80, for quantification by flow cytometry (gating strategies were shown in **Supplementary Fig. 9a**). As expected, in dexamethasone-treated mice, coupled processes of thymocyte apoptosis and phagocytic clearance of dead cells led to reduced thymus weight (**Fig. 5b**) and the total number of cells per thymus (**Fig. 5c**), accompanied by increased macrophage infiltration (**Fig. 5d**) and a remarkably higher percentage of Annexin V^+^ cells in the thymus (**Fig. 5e**). We did not observe a significant change in thymus weight or the total number of cells in *Wdfy3* knockout mice compared to controls, yet myeloid *Wdfy3*-deficiency led to an increased percentage of Annexin V^+^ cells, implying impaired efferocytic clearance of apoptotic thymocytes (**Fig. 5e**). Note that the impaired efferocytic clearance in Cre^+^ mice was unlikely to be caused by reduced macrophage availability because the percentage of macrophages per thymus was comparable between Cre^−^ and Cre^+^ mice treated with either PBS or dexamethasone (**Fig. 5d**). To assess efferocytosis in the thymus in situ, thymus sections were labeled and fluorescently imaged for TUNEL^+^ cells (ACs) that were either associated with CD68^+^ macrophages as a result of efferocytosis, or not associated with macrophages, i.e. “free” ACs, indicating inefficient efferocytic clearance. The ratio of macrophage-associated vs. free ACs was significantly lower in Cre^+^ mice, further supporting impaired efferocytosis in *Wdfy3* knockout mice (**Fig. 5f**).

For PM efferocytosis (**Fig. 5g**), TAMRA-labeled apoptotic thymocytes were injected intraperitoneally into Cre^−^ and Cre^+^ mice. 15 minutes after injection, peritoneal exudate was collected and stained for F4/80 to identify macrophages. The percentage of TAMRA^+^ PMs was quantified by flow cytometry (gating strategies were shown in **Supplementary Fig. 9b**). Consistent with thymus efferocytosis, the percentage of TAMRA^+^ PMs was significantly lower in Cre^+^ mice (**Fig. 5h**), supporting reduced AC efferocytosis by PMs *in vivo*.

### WDFY3 is required for efferocytosis in human macrophages

We further confirmed that in human macrophages, knockdown of WDFY3 by transfection of small interfering RNA (siRNA) led to impaired uptake and degradation of engulfed ACs during efferocytosis (**Fig. 6a-6e**). Human CD14^+^ monocytes were isolated from buffy coats of three independent subjects and differentiated to macrophages (human monocyte-derived macrophages, HMDMs) using M-CSF. On day 5, non-targeting control siRNA pool or WDFY3-targeting siRNA pool were transfected using Lipofectamine RNAiMAX (**Fig. 6a**). At 48-hour post-transfection, efficient knockdown of WDFY3 was confirmed at both mRNA (**Fig. 6b**) and protein levels (**Fig. 6c**). We then performed efferocytosis of human Jurkat cells labeled by both Hoechst and pHrodo. Consistent with the results in murine macrophages, both uptake and acidification (**Fig. 6d**) of ACs were impaired in HMDMs with siRNA-mediated *WDFY3* knockdown. The percentage of HMDMs with non-fragmented ACs was greater with *WDFY3* knockdown (**Fig. 6e**), confirming impaired degradation. The results support the role of WDFY3 in efferocytosis by human primary macrophages. This result is consistent with WDFY3’s level of conservation across mammalian species.

**Fig. 6.**
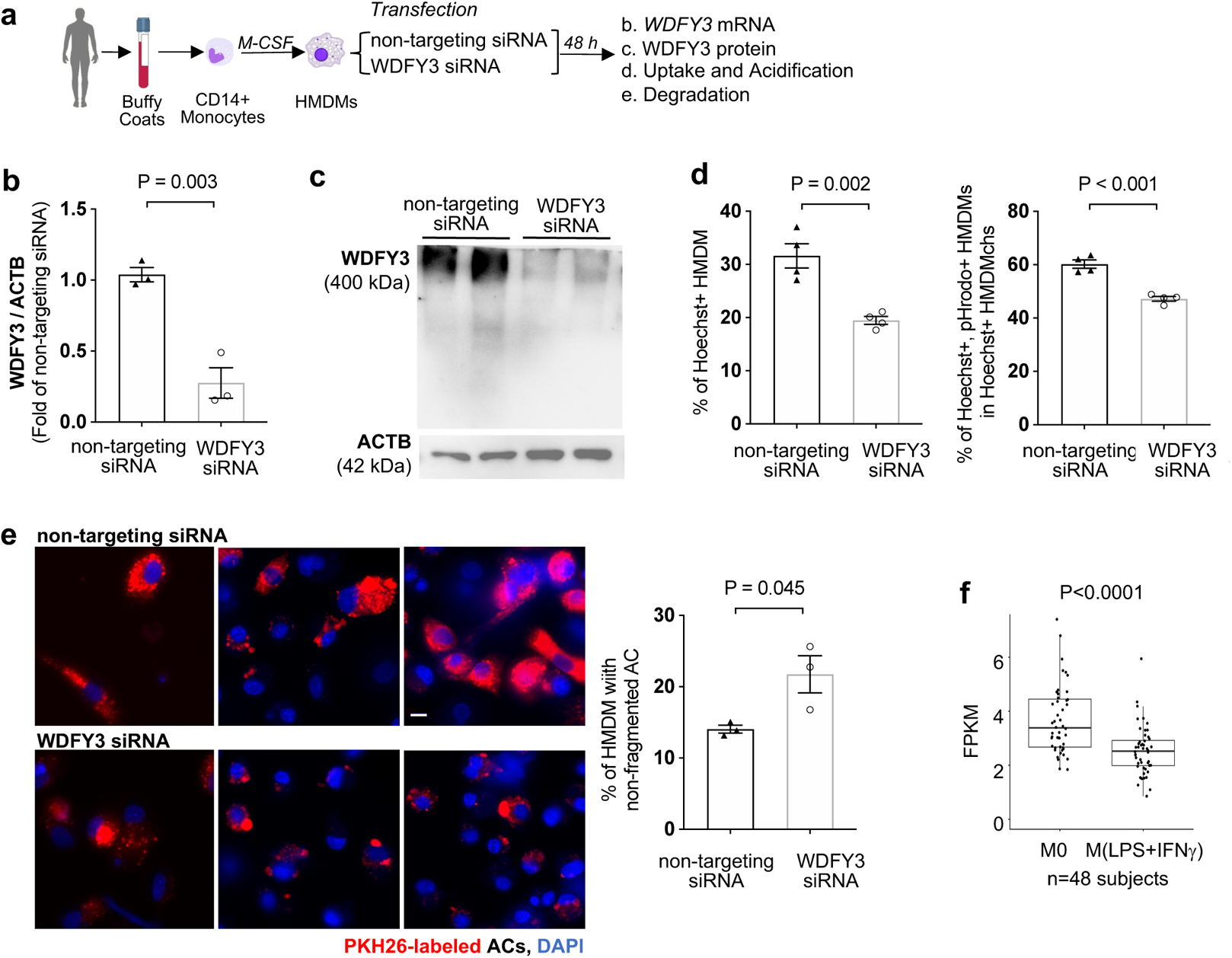
WDFY3 regulates efferocytosis in human macrophages. **(a)** Schematics of human monocyte differentiation to macrophages (HMDMs) and knockdown of WDFY3 with Lipofectamine RNAiMAX-mediated transfection of siRNAs targeting WDFY3, or non-targeting siRNAs as the control. **(b)** Validation of knockdown efficiency at mRNA level by qRT-PCR (n = 3 biological replicates, each from the average of 3 technical replicates). **(c)** Validation of knockdown efficiency at protein level by western blot (n = 2 biological replicates, data are representative of three independent experiments). **(d)** Efferocytosis of apoptotic Jurkat cells labeled by both Hoechst and pHrodo. The percentage of HMDMs with Hoechst-labeled ACs (indicating uptake), and the percentage of Hoechst^+^/pHrodo^+^ HMDMs in Hoechst^+^ HMDMs (indicating acidification upon uptake) were quantified by flow cytometry. Both uptake and acidification of ACs were impaired in HMDMs with siRNA-mediated WDFY3 knockdown (n = 3 biological replicates, each from the average of 2 technical replicates). **(e)** Fragmentation of engulfed ACs was assessed three hours after washing away the unengulfed ACs. The percentage of HMDMs with non-fragmented PKH26 staining in all PKH26^+^ HMDMs was determined (n = 3 biological replicates). **(f)** RNA-seq was performed for HMDMs either unstimulated (M0) or treated with LPS and IFNγ for 18-20 hours (M1-like). The expression of *WDFY3* was visualized (n = 48 biological replicates).

We also set out to determine how WDFY3 may be regulated during inflammation using our previously published RNA-seq data of human HMDMs, either unstimulated (M0) or stimulated with LPS and IFNγ (M1-like) for 18-20h^42^. M1-like inflammatory stimulation reduced *WDFY3* mRNA (**Fig. 6f**). Thus, reduced *WDFY3* during inflammation and subsequently impaired WDFY3-mediated macrophage efferocytosis may contribute to impaired efferocytic clearance of ACs *in vitro* and *in vivo*, exacerbating inflammation.

## Discussion

We developed a genome-wide CRISPR screen in primary macrophages. By focusing on efferocytosis, a complex macrophage functional phenotype, we illustrated the versatility of pooled screens and provided an effective approach for genome-wide CRISPR screening in primary macrophages derived from Cas9 transgenic mice. We have identified many known genes regulating efferocytosis and general phagocytosis, illuminating the most important genes essential for the uptake of ACs during efferocytosis. We have also uncovered and validated WDFY3 as a novel regulator specifically regulating the phagocytosis of dying cells, but not other substrates, using orthogonal assays *in vitro* and *in vivo*. Mechanistically, WDFY3 deficiency led to impaired phagosome formation due to defects in actin depolymerization. We further revealed that WDFY3 directly interacts with GABARAP, one of the seven members of the LC3/GABARAP protein family, to facilitate LC3 lipidation and the efficient degradation of the engulfed cargo (**Fig. 7** for a schematic summary). Further, WDFY3 expression was suppressed by inflammatory stimulation. Thus, WDFY3 regulates multiple steps during efferocytosis. Targeting WDFY3 may have therapeutic implications for diseases related to defective efferocytosis.

**Fig. 7.**
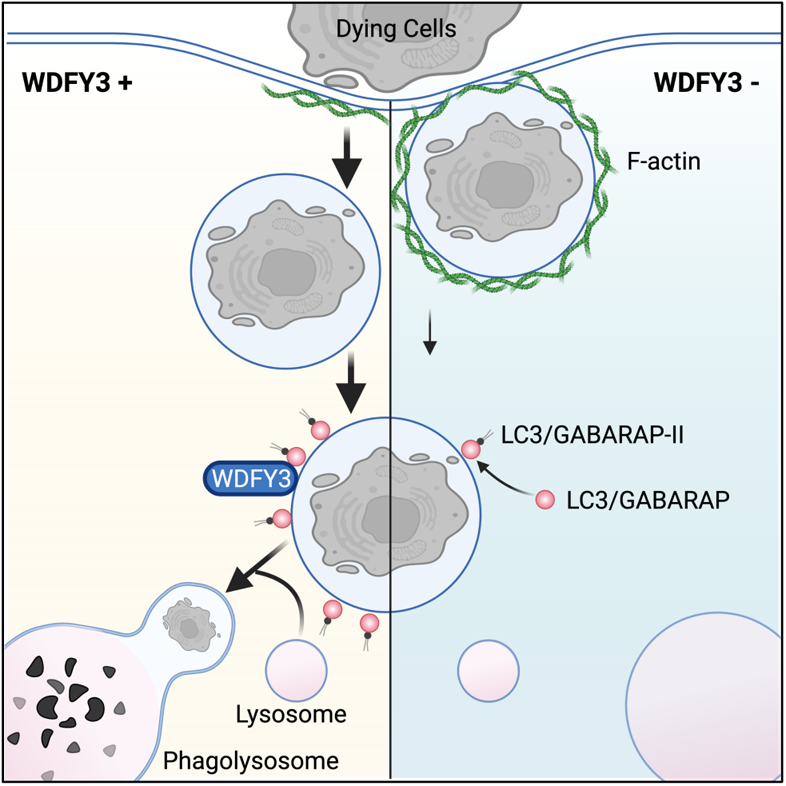
Schematic figure summarizing how WDFY3 regulates efferocytosis by macrophages. WDFY3 was discovered as a new regulator of efferocytosis by macrophages. WDFY3 deficiency in macrophages specifically impaired uptake, not binding, of apoptotic cells due to defective actin depolymerization, thus phagosome formation. WDFY3 directly interacts with GABARAP, one of the seven members of the LC3/GABARAP protein family, to facilitate LC3 lipidation and the subsequent phagosome lysosome fusion and degradation of the engulfed AC components.

We unexpectedly uncovered a novel role of WDFY3 in LAP. The detailed molecular mechanisms require further investigation. FYVE domain is known to bind phosphatidylinositol 3-phosphate (PI3P). PI3P is produced during both autophagosome and phagosome formation and is required for the recruitment of autophagic machinery for downstream fusion with lysosomes. It is therefore plausible that during efferocytosis, WDFY3 may be recruited through its known PI3P binding domain and acts as a scaffold that bridges ACs and autophagic machinery to regulate phagosome-lysosome fusion and lysosomal degradation of the engulfed cargos. Other questions remain unanswered, e.g. whether identical or different functional domains and binding partners have been involved in efferocytosis vs. in WDFY3-mediated aggrephagy; what is the function of N-terminal WDFY3; which molecular domains are required and sufficient for the role of WDFY3 in uptake and/or degradation and what are their protein-protein interaction partners. Pull-down experiments using specific domains of WDFY3 followed by quantitative proteomic screening, and live-cell imaging of endogenously tagged WDFY3 at baseline and during efferocytosis will further uncover these molecular mechanisms.

WDFY3 was highly expressed in myeloid cells compared with other immune cells. WDFY3 expression in HMDM was reduced by proinflammatory stimulation with LPS and IFNγ. The role of WDFY3-mediated efferocytosis in inflammation resolution and the therapeutic potential to enhance WDFY3 in diseases related to defective efferocytosis warrant further investigation. Indeed, overexpression of C-terminal WDFY3 (WDFY3_2981-3526_) can enhance aggrephagy in neurons as indicated by increased aggregate clearance^33, 41^, supporting the therapeutic premise to target macrophage WDFY3 to stimulate efferocytosis. Therapeutic activation of WDFY3 may represent a pro-efferocytotic therapy in atherosclerosis and other diseases related to defective efferocytosis.

The screen has implied many highly ranked, potentially novel regulators of macrophage efferocytosis. Among the top-ranked positive regulators, in addition to *Wdfy3*, *Sh3glb1, Snx24*, and *Vps33a* are annotated in autophagy-related pathways (**Supplementary Table 3**). The generation of PI3P on the phagosomal membrane recruits LC3-conjugation machinery, and abrogation of LC3 lipidation at the membrane impairs phagosome maturation and lysosome-mediated degradation^1, 28^. *Sh3glb1* activates lipid kinase activity of PIK3C3 during autophagy by associating with the PI3K complex II (PI3KC3-C2)^43^. *Snx24* contains a PX domain that mediates specific binding to membranes enriched in PI3P^44^. *Vps33a* is required for lysosome fusion with endosomes and autophagosomes^45^. These top screen hits may represent additional novel components of the cellular machinery that regulates efferocytosis. These promising targets and other potential novel regulators as uncovered by the screen have tremendous potential for additional novel discoveries. In our genome-wide screen, we employed a strategy of interpreting the results with a relatively permissive FDR threshold. Secondary screens with an increased number of gRNAs per gene and the number of cells infected per gRNA are expected to further improve the specificity and sensitivity for pooled screens in primary cells^46^.

Furthermore, this screening platform can be adapted to screen for phagocytic regulators of distinct substrates, e.g. bacteria and amyloid-β aggregates, for which the engulfment by physiologically-relevant primary macrophages will be more informative, and to study gene pairs with epistatic interactions using libraries with multiplexed gRNAs. Our platform will facilitate the identification of efferocytosis regulators affecting distinct molecular steps, including recognition and degradation. For example, by applying different selection strategies to separate macrophages with engulfed and acidified cargos from those with engulfed yet non-acidified cargos, genes specifically regulating the intracellular processing and degradation can be systematically interrogated. Further, screening for regulators responsible for efferocytosis of dying cells undergoing different modes of cell death can be studied. Since the number of macrophages required for genome-wide coverage and the required Cas9 transgenic expression makes it impractical for genome-wide pooled screens to be performed in human primary macrophages, screens in primary murine macrophages provide opportunities for physiologically-relevant discoveries of novel biology, which can then be validated in human macrophages. Our experimental framework also provides a general strategy for systematic identification of genes of interest and uncovering novel regulators of complex macrophage functions, such as lipid uptake and foam cell formation. This genetic platform promises to accelerate clinically relevant, mechanism-based translational research projects in macrophage biology and related human diseases.

**In summary,** we have established a pooled genome-wide CRISPR knockout screen in primary macrophages for discoveries of novel regulators of macrophage efferocytosis. The screen has revealed WDFY3 as a novel regulator of efferocytosis *in vitro* and *in vivo*, in the mouse and in human cells. The findings advanced our understanding of fundamental mechanisms of efferocytosis regulated by WDFY3. The screen top hits may likely contain additional novel regulators that can be further validated and promise to yield insights into diseases manifested by dysregulated efferocytosis. The innovative screen approaches established in this project are of broad and fundamental value to the community for conducting functional screens of novel regulators of complex macrophage function.

## Methods

The source of cell lines, mouse strains, gRNA sequences, siRNAs, primers, plasmids, antibodies, chemicals, and other assay kits and reagents were summarized in the Supplemental Information.

### Cell Lines

Cell lines, including Jurkat (lymphocytes, human acute T cell leukemia), THP-1 (monocytes, human acute monocytic leukemia), U937 (monocytes, human histiocytic lymphoma), and L-929 (mouse fibroblasts) were obtained from ATCC and handled according to the instructions provided on the ATCC product sheet.

### Bone Marrow Isolation and Differentiation to Bone Marrow-Derived Macrophages (BMDMs)

Bone marrow (BM) cells from 8-12 week old mice were isolated by flushing femurs and tibia with DMEM basal medium using 10 mL syringes with 22G needles. The isolated BM cells were cultured at 37°C, 5% CO_2_ on non-tissue-culture-treated vessels for 7-10 days in BMDM culture medium containing DMEM supplemented with 10% (vol/vol) heat-activated fetal bovine serum (HI-FBS), 20% (vol/vol) L-929 fibroblast conditioned medium, and 2 mM L-Glutamine. During differentiation, the growth medium was replaced with fresh medium 96 h after seeding and then every 2-3 days. In vitro assays were performed in BMDMs from day 8 to day 10.

### Peritoneal Macrophage (PM) Isolation

Cold PBS was injected into the peritoneum of donor mice for a 10-min incubation. Peritoneal exudates were then collected using 10 mL syringes with 21G needles and plated on non-tissue-culture-treated vessels. Floating cells were removed 6 h after plating and the attached cells were used as resident peritoneal macrophages (PMs). PMs were maintained in DMEM supplemented with 10% (vol/vol) HI-FBS, 20% (vol/vol) L-929 fibroblast conditioned medium, and 2 mM L-Glutamine for 24 h at 37°C, 5% CO_2_ before indicated assays^47^.

### Human Monocyte Derived Macrophages (HMDMs)

Buffy coats of anonymous, de-identified healthy adults were obtained from the New York Blood Center (NYBC) for isolating peripheral blood mononuclear cells (PBMCs). Buffy coats were diluted with 1X DPBS supplemented with 2 mM EDTA at a 1:1 ratio, i.e. 8 mL buffy coats were diluted with 8 mL DPBS to a total volume of 16 mL. The diluted buffy coats were carefully laid on 9 mL Ficoll-Paque solution, i.e. a 4:3 ratio in 50-mL conical tubes and centrifuged at 400 x g for 40 min at 20°C without brake. PBMC layer was transferred and washed with washing buffer (1X DPBS, 2% (vol/vol) HI-FBS, 5 mM EDTA, 20 mM HEPES and 1 mM sodium pyruvate), centrifuged at 500 x g for 10 min at 4°C. The PBMC pellets were washed again in RPMI-1640 medium containing 20% (vol/vol) HI-FBS. The pellets were then resuspended and cultured in RPMI-1640 medium supplemented with 20% (vol/vol) HI-FBS and 100 ng/mL human macrophage colony-stimulating factor (M-CSF) for 7-10 days^48^. The growth medium was replaced with fresh medium 96 h after seeding and then every 2-3 days.

### THP-1 and U937 Differentiation to Macrophages

THP-1 human acute monocytic leukemia cell line was obtained from ATCC and grown in suspension in THP-1 culture medium containing RPMI-1640 supplemented with 10% (vol/vol) HI-FBS, 1 mM Sodium Pyruvate, 10 mM HEPES, and 50 μM 2-Mercaptoethanol. THP-1 macrophages were differentiated from THP-1 cells in the above culture media supplemented with 100 nM Phorbol 12-myristate 13-acetate (PMA) for 24 h at a seeding density of 1 × 10^6^ cells per well of a 6-well tissue culture plate. PMA-containing media was then removed and replaced with THP-1 culture media for 48 h culture. The same seeding density was used for U937 differentiation to macrophages with 50 nM PMA for 3 days^11^.

### Experimental Animals

Animal protocols were approved by Columbia University’s Institutional animal care and use committee. All animals were cared for according to the NIH guidelines. Mice were socially housed in standard cages at 22°C under a 12-12 h light-dark cycle with *ad libitum* access to water and food provided by the mouse barrier facility. Rosa26-Cas9 knockin mice were obtained from the Jackson Laboratory (Cat# 026179) (female mice were used for the CRISPR screen and validation). *Wdfy3^fl/fl^* mice were obtained from Dr. Ai Yamamoto’s lab^34^ and Dr. Konstantinos Zarbalis’s lab^26^. Myeloid-specific Wdfy3 knockout mice were created by crossing LysMCre^+/−^ mice with *Wdfy3^fl/fl^* mice. LysMCre^+/−^*Wdfy3^fl/fl^* mice (Cre^+^) had myeloid-specific knockout of *Wdfy3*, while LysMCre^−/−^*Wdfy3^fl/fl^* littermates (Cre^−^) served as controls (both male and female mice were used).

### Lentiviral Plasmid Construction

The Brie murine CRISPR knockout pooled library in the lentiGuide-Puro backbone was obtained from Addgene (#73663)^17^. To validate the top screen hits using individual gRNAs, pairs of oligonucleotides with BsmBI-compatible overhangs were separately annealed and cloned into the lentiGuide-Puro vector (Addgene #52963) using standard protocols available via https://www.addgene.org/52963/. To validate the role of *Wdfy3* using a separate plasmid platform, gRNA targeting *Wdfy3* was selected from the murine Sanger lentiviral CRISPR library (Sigma) and the *Wdfy3*-targeting lentiviral vector, as well as the non-targeting control vector, were obtained (Sigma). To overexpress C-terminal WDFY3, pLE4-WDFY3_2543-3526_ was constructed by inserting Myc-WDFY3_2543-3526_, which was from pcDNA-myc-WDFY3_2543-3526_ provided by Dr. Ai Yamamoto^33^, into the pLE4 lentiviral backbone. eGFP was then inserted into the N-terminal of WDFY3 to generate pLE4-eGFP-WDFY3_2543-3526_ to allow the identification of WDFY3-overexpressing population by flow cytometry upon transduction.

### Lentiviral Packaging and Transduction

Lentivirus particles were generated from HEK293T cells (ATCC CRL-3216) by co-transfection of lentiviral vectors with the packaging plasmid psPAX2 (Addgene #12260) and envelope plasmid pMD2G (Addgene #12259) using FuGene 6 transfection reagent (Promega). The medium was changed 18 h after transfection. 48 h after transfection, lentiviral supernatants were harvested and filtered through 0.45-µm SFCA filters (Corning). Lentiviral particles were further concentrated using Lenti-X concentrator (Clontech) following the manufacturer’s instructions.

Mouse BM cells were isolated and plated (day 0). On day 1, BM cells were virally transduced in BMDM culture medium supplemented with 10 μg/mL polybrene. On day 2, half of the medium was replenished with fresh BMDM culture medium. On day 6, the transduced cells underwent puromycin selection at 5 µg/mL for 48 h. On day 9, i.e. 24h after removing puromycin, BMDMs were used for efferocytosis assays. The pLE4 lentiviral vector does not have a puromycin-resistant gene, thus no antibiotics selection was performed.

### Induction of Apoptosis and Fluorescent Labeling of Apoptotic Cells (ACs)

Apoptotic Jurkat cells were generated by treating Jurkat cells with 5 μg/mL staurosporine in RPMI-1640 medium for 3 hours at a density of 2.5 × 10^6^ cells/mL at 37°C, 5% CO_2_. The method routinely yields greater than 90% Annexin V-positive apoptotic Jurkat cells. After washing in 1X DPBS, apoptotic Jurkat cells were resuspended at a concentration of 2 × 10^7^ cells/mL in Diluent C with either PKH67 (green fluorescence) or PKH26 (red fluorescence) per the manufacturer’s instruction. After labeling, the cells were rinsed twice with DMEM basal medium containing 10% HI-FBS and immediately used for efferocytosis assay. For labeling with other fluorescent probes, ACs were resuspended at a density of 2.5 × 10^6^ cells/mL and incubated with 20 ng/mL pHrodo red (Life Technologies) and Hoechst 33342 (Invitrogen) for 30 min in 1X DPBS.

To isolate mouse thymocytes and induce apoptosis, thymi were dissected from C57BL/6J mice and were grounded and filtered through a 40 μm cell strainer to obtain single-cell suspension. The induction of apoptosis was initiated by incubating the thymocytes with 50 µM dexamethasone in DMEM at 37 °C, 5% CO_2_ for 4 h. The cells were then stained with 10 μg/mL TAMRA (Invitrogen) at a concentration of 2 × 10^7^ cells/mL in serum-free DMEM for 25 min.

### Preparation of Sheep Red Blood Cells (RBCs) for Efferocytosis

Sheep red blood cells (RBCs) (Rockland) were obtained. For heat-shock treatment, RBCs were incubated under 56°C in a water bath for 5 min^49^. For IgG-opsonization, RBCs were incubated with 1 μg/mL anti-RBCs antibodies in DMEM basal medium containing 10% (vol/vol) HI-FBS to conjugate with IgG at 37°C, 5% CO_2_ for 1.5 h^49^. The non-treated, heat-shock treated or IgG-conjugated RBCs were labeled with PKH67 following the same procedures for the labeling of apoptotic Jurkat cells.

### In Vitro Efferocytosis and Phagocytosis Assays

For imaging-based quantification, macrophages were plated in 96-well plates at a density of 0.3 × 10^5^ per well. For flow cytometry-based quantification, macrophages (BMDMs, PMs, or HMDMs) were plated in 6-well or 24-well plates at a density of 1.5 × 10^6^ per well or 0.2 × 10^6^ per well, respectively. Fluorescently-labeled apoptotic cells were co-incubated with macrophages at a 5:1 AC : macrophage ratio for 1 h (or as described in Figures) at 37°C, 5% CO_2_. Macrophages were then washed with 1X DPBS gently to remove unbound targets. For imaging-based quantification, macrophages were fixed with 2% PFA for 30 min, rinsed 3 times with 1X DPBS, and counterstained by DAPI. For flow cytometry-based quantification, macrophages were lifted using CellStripper, a non-enzymatic cell dissociation solution, for live cell analysis. The phagocytosis of beads, RBCs, and zymosan particles by BMDMs was determined upon incubation for 1 h at the specific ratio or concentration as specified in the respective figures.

### CRISPR-Cas9 Screen for efferocytosis in BMDMs

CRISPR-Cas9 screens were performed using the Brie library^17^. BM cells isolated from *Rosa-Cas9* knockin mice were virally transduced at a low multiplicity of infection (MOI) of 0.3 and targeting ~1,000 fold coverage of the library. After puromycin selection, BMDMs were dissociated and replated in 10-cm tissue culture plates at a density of 6 × 10^6^ per plate for two-round efferocytosis. For the 1^st^ round, PKH67-labeled ACs were incubated with BMDMs at a 5:1 ratio for 45 min. After removing the unbound ACs, macrophages were rested for 3 h before the 2^nd^ round, in which PKH26-labeled ACs were incubated with BMDMs at a 5:1 ratio for 90 min. Unbound ACs were removed and BMDMs were collected for sorting on BD Influx. The sorted populations were processed individually for genomic DNA extraction using DNeasy Blood and Tissue Kit (Qiagen) and subjected to PCR reactions to generate the libraries. The purified PCR products were sequenced on Illumina NextSeq500 system to determine gRNA abundance in two independent replicates. Data were analyzed using MAGeCK (Model-based Analysis of Genome-wide CRISPR-Cas9 Knockout^21^ to obtain ranked lists of screen hits. Independent validation of top screen hits by individual gRNAs was performed by lentiviral transduction of gRNA in Rosa-Cas9 knockin BM cells and differentiation to BMDM followed by efferocytosis assays and quantification^23^.

### Analysis of Macrophage Capability of Binding

BMDMs were stained with 0.5 μM CellTracker Green CMFDA (5-chloromethylfluorescein diacetate) for 60 min. The CellTracker dye freely passes through cell membranes and is well-retained in cells, allowing labeling of cytoplasmic area. BMDMs were then treated with 5 μM cytochalasin D for 30 min. Cytochalasin D blocks the assembly and disassembly of actin monomers, thus preventing internalization of ACs. The treated BMDMs were then incubated with TAMRA-stained apoptotic mouse thymocytes for 30 min at 37°C, 5% CO_2_ to allow binding. The unbound ACs were extensively washed with 1X DPBS, BMDMs were fixed with 2% PFA for 30 min and washed with 1X DPBS for 3 times, followed by imaging with ImageXpress Micro 4 High-Content Imaging System with a Nikon Plan Apo λ 20x/0.75 objective lens to analyze the percentage of macrophages with bound ACs.

### Time-lapse Imaging of Phagosome Formation

BMDMs cultured on chambered coverslips with 8 individual wells (ibidi) at a density of 0.12 × 10^6^ were stained with 0.5 µM CellTracker Green CMFDA Dye (Invitrogen) in DMEM and 10% (vol/vol) HI-FBS for 60 min. The medium was replaced with fresh DMEM containing 10% HI-FBS and apoptotic Jurkat cells were added at a 5:1 AC : BMDM ratio. BMDMs were imaged with Nikon Ti Eclipse inverted microscope for spinning-disk confocal microscopy equipped with a 60x/1.49 Apo TIRF oil immersion lens. Images of the same fields were recorded at 30 s intervals for 20 min.

### Visualization and Quantification of F-actin Dynamics during Efferocytosis

BMDMs plated on 96-well plates were stained with 0.5 µM CellTracker Green CMFDA Dye (Invitrogen) and 1 µM SiR-actin (Cytoskeleton) for 60 min. ACs labeled by NCS-nucleomask blue (Invitrogen) were added to the macrophages to replace the staining medium at a 5:1 AC : macrophage ratio for 1 h efferocytosis. Macrophage monolayer was then vigorously washed with 1X DPBS to remove unbound ACs, fixed with 2% PFA for 30 min and washed with 1X DPBS for 3 times, and imaged by ImageXpress Micro4 high content microscopy (Molecular Device) with a Nikon Plan Apo λ 40X/0.95 objective lens. The percentage of macrophage with bright F-actin ring, as an indicator of F-actin polymerization, was quantified.

To quantify F-actin intensity by flow cytometry, BMDMs plated on 6-well non-tissue culture plates were incubated with Hoechst-labeled ACs for 1h. Unbound ACs were washed away and BMDMs were collected and fixed by 2% PFA for staining with 1 µM siR-actin in washing buffer (1X DPBS, 2% (vol/vol) HI-FBS, 5 mM EDTA, 20 mM HEPES and 1 mM sodium pyruvate). siR-actin-labeled F-actin levels were quantified as the mean fluorescence intensity (MFI) of siR-actin in BMDMs with or without engulfment of ACs.

### Analysis of Fragmentation of Engulfed AC Components

PKH26-labeled ACs were added to BMDMs or HMDMs and incubated for 45 min. Unengulfed ACs were removed by vigorous rinsing with 1X DPBS. After being cultured for an additional 3 hours, the macrophages were fixed with 2% PFA and counterstained with DAPI. Images were captured using ImageXpress Micro4 high content microscopy (Molecular Device) with a Nikon Plan Apo λ 40X/0.95 objective lens. The percentage of macrophages containing non-fragmented AC-derived fluorescence, which is a measure of AC corpse degradation, was quantified^50^.

### Membrane-bound LC3 Detection Assay

BMDMs were incubated with Hoechst-labeled ACs at a 5:1 AC : BMDM ratio at 37°C, 5% CO_2_ for 1 h efferocytosis. Unbounded ACs were washed away. BMDMs were collected and resuspended in 300 mL cold PBS with 20 µg/mL digitonin, and incubated on ice for 10 min to permeabilize cells and allow non-membrane bound LC3 to be removed from cells. Permeabilized BMDMs were then centrifuged for 5 min at 750 × g, followed by incubation with anti-LC3A/B-AF488 antibody diluted 1:500 in cold washing buffer (1X DPBS, 2% (vol/vol) HI-FBS, 5 mM EDTA, 20 mM HEPES and 1 mM sodium pyruvate) for 15 min on ice to stain the membrane-bound lipidated LC3-II within the cells. After staining, macrophages were washed with 1 mL cold washing buffer and were centrifuged for 5 min at 750 × g. Cell pellets were resuspended in washing buffer and acquired on a flow cytometer^40^.

### RNA Sequencing and Gene Set Enrichment Analysis

Total RNAs were extracted from BMDMs (male mice: 4 Cre^+^ and 4 Cre^−^) using the Quick-RNA miniprep plus kit (Zymo). With a minimum of 300 ng input RNA, strand-specific, poly(A)+ libraries were prepared and sequenced at 20 million 100-bp paired-end reads per sample. Raw sequencing reads were mapped to the mouse genome version GRCm39 (M27) using Salmon^51^ (version 1.5.1) to obtain transcript abundance counts. MultiQC was used to generate quality control reports based on Salmon read mapping results. The transcript-level count information was summarized to the gene level using tximport^52^ (version 1.20.0). Differential expression was assessed using DESeq2^53^ (version 1.34.0). Genes with an absolute fold change > 1.5 and false discovery rate (FDR)-adjusted P value < 0.05 were considered as differentially expressed (DE). The output of DESeq2 were scored and ranked based on P value and shrunken log_2_ fold change by apeglm^54^ using ranking metrics “−log10 P value multiplied by the sign of log-transformed fold-change”^55^. The ranked gene list was then used for Gene Set Enrichment Analysis (GSEA)^56^ (version 4.2.0) to identify the gene sets overrepresented at the top or bottom of the ranked list using the Human Reactome Pathway (the most actively updated general-purpose public database of human pathways) and the Gene Ontology Biological Process annotation (the most commonly used resource for pathway enrichment analysis) within the Molecular Signatures Database.

### Ingenuity Pathway Analysis

Ingenuity pathway analysis (IPA) software using build-in scientific literature based database (according to IPA Ingenuity Web Site, www.ingenuity.com) was used to identify canonical pathways, overrepresented in top-scored CRISPR screen hits.

### Quantitative RT-PCR

Total RNA was extracted using Quick-RNA Miniprep Kit (Zymo) and cDNA was synthesized using High-Capacity cDNA Reverse Transcription Kit (Applied Biosystems) as per the manufacturer’s instructions. To measure gene expression, quantitative RT-PCR was performed using POWERUP SYBR Green Master Mix by QuantStudio™ 7 Flex Real-Time PCR System (Applied Biosystem, 4485701). ΔΔCT method was used to analyze the relative levels of each transcript normalized to human *ACTB*.

### Immunoblotting and Immunoprecipitation

Macrophages were harvested and lysed in RIPA lysis buffer (Millipore) supplemented with protease inhibitor cocktail (Roche) and phosphatase inhibitor cocktail (Roche). Protein concentration was quantified using Pierce BCA protein assay kit (Thermo Fisher). Equal amount of protein were mixed with 5X SDS sample buffer [5%(vol/vol) β-Mercaptoethanol, 0.02%(vol/vol) Bromophenol blue, 30%(vol/vol) Glycerol, 10%(vol/vol) Sodium dodecyl sulfate, 250 mM Tris-Cl, pH 6.8)] (or 4X Bolt LDS sample buffer) and loaded onto a 3-8% Tris-Acetate NuPage gel (for WDFY3) or 10-20% Tris-glycine gel (for LC3) and then electro-transferred to a 0.45 μm (or 0.2 μm) PVDF membrane (Thermo Scientific). After blocking with 5% milk, the membrane was incubated with the indicated primary antibody overnight at 4°C. The membrane was then washed for 3 times in TBST and incubated with HRP-conjugated goat anti-rabbit IgG (1:5000 dilution) for 1 h at room temperature. After the final wash to remove unbound antibodies, the protein expression was detected by SuperSignal^TM^ West Pico PLUS Chemiluminescent Substrate (Thermo Scientific) and imaged using ChemiDoc Imaging System (Bio-rad). Band intensity was quantified using the software FIJI.

For immunoprecipitation, 100 μg total cell lysates were incubated with 4 μg anti-GABARAP antibodies in 500 RIPA buffer. A pool of 2 μg Rabbit anti-GABARAP (Cell Signaling Technology 13733S) and 2 μg Rabbit anti-GABARAP (N-term) (Abgent AP1821a) targeting different regions of the GABARAP protein were used to improve pull-down efficiency. The lysate and antibody mix was incubated overnight at 4°C, followed by a 1 h-incubation with 100 μL protein A/G agarose beads (Thermo Scientific Pierce) at 4°C. Immunoprecipitants were washed 3 times with lysis buffer and eluted with 4X LDS sample buffer (Invitrogen) by boiling at 70°C for 10 min. Immunocomplexes were subjected to 3-8% Tris-glycine gel and immunoblotting analysis for WDFY3.

### In Vivo Thymus Efferocytosis Assay

Cre^+^ and Cre^−^ mice of 8-12 weeks old were injected intraperitoneally with 200 μL PBS or 200 μL PBS containing 250 μg dexamethasone. Dexamethasone was prepared freshly by diluting 4X stock in DMSO with sterile PBS. 18 h after injection, mice were weighed and euthanized, and thymi were harvested and both lobes were weighed. One lobe was immersed in OCT and snap-frozen for immunohistochemical staining to determine efferocytosis *in situ*, while the other lobe was mechanically disaggregated into single-cell suspension for flow cytometry^50^.

To evaluate *in situ* efferocytosis^50^, frozen thymus specimens were cryosectioned at 4-µm and placed on Superfrost plus microscope slides. Sections were fixed in 4% PFA for 10 mins and permeabilized in 1% Triton X-100 for 15 mins. After rinsing with PBS for three times, sections were incubated with TUNEL staining reagents at 37°C for 60 min and then washed three times with PBS. Sections were then blocked with 5% goat serum for 60 min at room temperature, followed by overnight incubation at 4°C in anti-CD68 antibodies (Abcam) diluted in PBS supplemented with 5% BSA to label macrophage area. After washing in PBS, sections were incubated with fluorescently-labeled secondary antibodies and counterstained with DAPI. Images were captured using ImageXpress Micro4 with a Nikon Plan Apo 40X/0.95 objective lens. For quantification, the TUNEL+ nuclei in close proximity or in contact with CD68+ macrophages were counted as macrophage-associated ACs, indicative of efferocytosis. The TUNEL+ nuclei without neighboring macrophages were counted as free ACs. The ratio of macrophage-associated ACs to free ACs was calculated to represent the capability of efferocytosis by thymus macrophages.

To evaluate the percentage of Annexin V^+^ ACs by flow cytometry, mechanically disaggregated thymus cells were rinsed twice with cold PBS containing 2% HI-FBS and 1 mM EDTA. Cells were then stained with AF647-conjugated Annexin V (Thermo Fisher) in Annexin V binding buffer (Invitrogen) at a concentration of 5 × 10^6^ cells/mL for 15 min at room temperature, followed by analysis flow cytometry.

### In Vivo Peritoneal Macrophage Efferocytosis Assay

Cre^+^ and Cre^−^ mice of 12 weeks old were injected intraperitoneally with 1 × 10^7^ TAMRA-stained apoptotic mouse thymocytes in 300 µl PBS. 15 min after injection, mice were euthanized and peritoneal exudates were collected. The pelleted cells were stained by FITC-conjugated F4/80 antibody (BioLegend) to label macrophages. The percentage of TAMRA+ PMs was determined by flow cytometry^47^.

### siRNA-Mediated Gene Silencing and Transfection

Non-targeting siRNA and WDFY3-targeting siRNA (Dharmacon) were transfected using Lipofectamine RNAiMAX (Invitrogen) as per the manufacturer’s recommendation. Briefly, human PBMCs were seeded at 4 × 10^5^ per well of 24-well culture dish for differentiation to HMDMs for 7 days with ~70% confluence. HMDMs were then transfected with a final concentration of 25 pmol siRNA and 1 μL Lipofectamine RNAiMAX in 500 μL Opti-MEM (Invitrogen) for 6 h. A second transfection with the same condition was performed 18 h after the completion of the first transfection. HMDMs were collected 48 h from the start of the first transfection for assessing mRNA and protein expression, and efferocytosis capacities.

### Mouse Complete Blood Cell Count (CBC) and Differential Count

Retro-orbital bleeding was performed to collect ~500 μL blood per mouse for complete blood count and differential count using a Heska Element HT5 by the diagnostic lab at the Institute of Comparative Medicine, Columbia University Irvine Medical Center.

### Statistical Analyses

Statistical analyses were performed using GraphPad Prism 7. Nonparametric tests were used when sample size (n) was less than or equal to 5. When n > 6, data were first tested for normality using the D’Agostino-Pearson test (when n >=8) or Shapiro-Wilk test (when n < 8). F-test of equality of variances was performed to compare the two sample variances. Data that passed normality tests were presented as mean ± standard error of mean (SEM) and analyzed using Student’s t-test for two groups with one variable tested and equal variances (or with Welch’s correction if F-test was not satisfied); one-way ANOVA with Tukey’s post-hoc analysis for more than two groups with one variable tested; or two-way ANOVA with Tukey’s post-hoc analysis for more than two groups with two independent variables tested. Data that were analyzed using nonparametric tests were presented as median ± 95% confidence interval. Statistical significance of difference was accepted when P values were < 0.05. The specific P values, the number of independent experiments or biological replicates, and the number of technical replicates per independent experiment and biological replicate were specified in figures and figure legends.

## Data availability

The authors declare that all data supporting the findings of this study are available within the paper and its supplementary information files. Source data are provided with this paper. The datasets generated during the current study, including RNA-seq and CRISPR screening sequencing data will be deposited in the Gene Expression Omnibus (GEO) upon acceptance of the manuscript. The human macrophage RNA-seq data was previously published and are available at DRYAD with identifier doi:<u>10.5061/dryad.866t1g1nb</u>.

## Code availability

All code for data analysis associated with the current submission will be available at https://github.com/hanruizhang/ upon acceptance of the manuscript. Any updates will also be available via the above GitHub repository.

## Acknowledgments

The authors’ research work has received funding from R00HL130574 and R01HL151611, and the Irving Scholar award through UL1TR001873 by the National Center for Advancing Translational Sciences (NCATS) and National Institutes of Health (NIH) (to HZ), R00HL145131 (to AYJ), R21MH115347 and grant by Shriners Hospitals for Children (to KSZ), R35HL145228 (to IT), R01NS077111 and R01NS101663 (to AY), the Russell Berrie Diabetes Foundation Diabetes Scholar Program (to XW), American Heart Association Postdoctoral Fellowships (to XW, FL), and an NSF predoctoral fellowship (to KRC). The content in this manuscript is solely the responsibility of the authors and does not necessarily represent the official views of the NIH.

We would like to acknowledge the NIH funding sources to the Columbia Center for Translational Immunology (CCTi) Flow Cytometry Core by grant number S10OD020056 and S10RR027050 and P30DK063608; the NIH-supported microscopy resources in the Center for Biologic Imaging, specifically the confocal microscope supported by grant number 1S10OD019973-01; the NIH/NCI Cancer Center Support Grant P30CA013696 for the use of resources at the Columbia Genome Center; the Columbia Stem Cell Initiative (CSCI) Flow Cytometry Core under the leadership of Michael Kissner; and the High-Throughput Screening Facility at the JP Sulzberger Columbia Genome Center under the leadership of Dr. Charles Karan.

FACS cell sorting was performed with great help from Dr. Caisheng Lu, the Technical Director of the CCTi Flow Cytometry Core. We thank Ms. Xiaoli Sky Wu and Dr. Kenneth Chang at the Cold Spring Harbor Laboratory for their technical inputs for CRISPR screening design. We thank Dr. Oren Parnas at the Hebrew University of Jerusalem for his inputs on genome-wide CRISPR screening in primary cells. We thank Dr. Young Joo Yang for technical advice on the characterization of WDFY3.

Schematic figures were created with BioRender.com.

## Author contributions

J.S., X.W., and H.Z. conceived and designed the study. J.S., X.W., Z.W., F.L., Y.M., R.M.M., J.C., H.Z. performed experiments. J.S. X.W., Z.W., H.Z. analyzed the data and prepared the figures. J.S. and H.Z. performed CRISPR screening, and the bioinformatic analyses of CRISPR screening and murine RNA-seq data. C.X. performed bioinformatic analyses of human macrophage RNA-seq data. J.S. and H.Z. wrote the paper with inputs from all authors. K.R.C., A.Y. Jr, J.G.D., W.L. provided guidance for key techniques. K. S. Z. and A.Y. provided key reagents, mice, and critical technical inputs. I.T., A.Y. advised on the project and critically reviewed the paper. H.Z. mentored the performance of the work and supervised the funding.

## References

1. Boada-Romero, E., Martinez, J., Heckmann, B. L. & Green, D. R. The clearance of dead cells by efferocytosis. Nat Rev Mol Cell Biol, doi:10.1038/s41580-020-0232-1 (2020).

2. Doran, A. C., Yurdagul, A., Jr. & Tabas, I. Efferocytosis in health and disease. Nature reviews. Immunology, doi:10.1038/s41577-019-0240-6 (2019).

3. Morioka, S., Maueroder, C. & Ravichandran, K. S. Living on the Edge: Efferocytosis at the Interface of Homeostasis and Pathology. Immunity 50, 1149–1162, doi:10.1016/j.immuni.2019.04.018 (2019).

4. Trzeciak, A., Wang, Y. T. & Perry, J. S. A. First we eat, then we do everything else: The dynamic metabolic regulation of efferocytosis. Cell metabolism, doi:10.1016/j.cmet.2021.08.001 (2021).

5. Greenlee-Wacker, M. C. Clearance of apoptotic neutrophils and resolution of inflammation. Immunological reviews 273, 357–370, doi:10.1111/imr.12453 (2016).

6. Kojima, Y., Weissman, I. L. & Leeper, N. J. The Role of Efferocytosis in Atherosclerosis. Circulation 135, 476–489, doi:10.1161/circulationaha.116.025684 (2017).

7. Yurdagul, A., Jr., Doran, A. C., Cai, B., Fredman, G. & Tabas, I. A. Mechanisms and Consequences of Defective Efferocytosis in Atherosclerosis. Frontiers in cardiovascular medicine 4, 86, doi:10.3389/fcvm.2017.00086 (2017).

8. Hayat, S. M. G. et al. CD47: role in the immune system and application to cancer therapy. Cell Oncol (Dordr) 43, 19–30, doi:10.1007/s13402-019-00469-5 (2020).

9. Silva, E., Au-Yeung, H. W., Van Goethem, E., Burden, J. & Franc, N. C. Requirement for a Drosophila E3-ubiquitin ligase in phagocytosis of apoptotic cells. Immunity 27, 585–596, doi:10.1016/j.immuni.2007.08.016 (2007).

10. Sedlyarov, V. et al. The Bicarbonate Transporter SLC4A7 Plays a Key Role in Macrophage Phagosome Acidification. Cell host & microbe 23, 766–774.e765, doi:10.1016/j.chom.2018.04.013 (2018).

11. Haney, M. S. et al. Identification of phagocytosis regulators using magnetic genome-wide CRISPR screens. Nature genetics 50, 1716–1727, doi:10.1038/s41588-018-0254-1 (2018).

12. Kamber, R. A. et al. Inter-cellular CRISPR screens reveal regulators of cancer cell phagocytosis. Nature 597, 549–554, doi:10.1038/s41586-021-03879-4 (2021).

13. Penberthy, K. K. & Ravichandran, K. S. Apoptotic cell recognition receptors and scavenger receptors. Immunological reviews 269, 44–59, doi:10.1111/imr.12376 (2016).

14. Schlam, D. et al. Phosphoinositide 3-kinase enables phagocytosis of large particles by terminating actin assembly through Rac/Cdc42 GTPase-activating proteins. Nature communications 6, 8623, doi:10.1038/ncomms9623 (2015).

15. Andreu, N. et al. Primary macrophages and J774 cells respond differently to infection with Mycobacterium tuberculosis. Scientific reports 7, 42225, doi:10.1038/srep42225 (2017).

16. Platt, R. J. et al. CRISPR-Cas9 knockin mice for genome editing and cancer modeling. Cell 159, 440–455, doi:10.1016/j.cell.2014.09.014 (2014).

17. Doench, J. G. et al. Optimized sgRNA design to maximize activity and minimize off-target effects of CRISPR-Cas9. Nature biotechnology 34, 184–191, doi:10.1038/nbt.3437 (2016).

18. Li, W. et al. MAGeCK enables robust identification of essential genes from genome-scale CRISPR/Cas9 knockout screens. Genome biology 15, 554, doi:10.1186/s13059-014-0554-4 (2014).

19. Li, W. et al. Quality control, modeling, and visualization of CRISPR screens with MAGeCK-VISPR. Genome biology 16, 281, doi:10.1186/s13059-015-0843-6 (2015).

20. Chen, C. H. et al. Improved design and analysis of CRISPR knockout screens. Bioinformatics (Oxford, England) 34, 4095–4101, doi:10.1093/bioinformatics/bty450 (2018).

21. Wang, B. et al. Integrative analysis of pooled CRISPR genetic screens using MAGeCKFlute. Nature protocols 14, 756–780, doi:10.1038/s41596-018-0113-7 (2019).

22. Zhao, D. et al. Frontline Science: Tim-3-mediated dysfunctional engulfment of apoptotic cells in SLE. J Leukoc Biol 102, 1313–1322, doi:10.1189/jlb.3HI0117-005RR (2017).

23. Nakahashi-Oda, C. et al. CD300a blockade enhances efferocytosis by infiltrating myeloid cells and ameliorates neuronal deficit after ischemic stroke. Sci Immunol 6, eabe7915, doi:10.1126/sciimmunol.abe7915 (2021).

24. Dragich, J. M. et al. Autophagy linked FYVE (Alfy/WDFY3) is required for establishing neuronal connectivity in the mammalian brain. eLife 5, doi:10.7554/eLife.14810 (2016).

25. Lerm, M., Brodin, V. P., Ruishalme, I., Stendahl, O. & Särndahl, E. Inactivation of Cdc42 is necessary for depolymerization of phagosomal F-actin and subsequent phagosomal maturation. Journal of immunology 178, 7357–7365, doi:10.4049/jimmunol.178.11.7357 (2007).

26. Orosco, L. A. et al. Loss of Wdfy3 in mice alters cerebral cortical neurogenesis reflecting aspects of the autism pathology. Nature communications 5, 4692, doi:10.1038/ncomms5692 (2014).

27. Martinez, J. et al. Molecular characterization of LC3-associated phagocytosis reveals distinct roles for Rubicon, NOX2 and autophagy proteins. Nature cell biology 17, 893–906, doi:10.1038/ncb3192 (2015).

28. Martinez, J. LAP it up, fuzz ball: a short history of LC3-associated phagocytosis. Current opinion in immunology 55, 54–61, doi:10.1016/j.coi.2018.09.011 (2018).

29. Cunha, L. D. et al. LC3-Associated Phagocytosis in Myeloid Cells Promotes Tumor Immune Tolerance. Cell 175, 429–441.e416, doi:10.1016/j.cell.2018.08.061 (2018).

30. Heckmann, B. L. & Green, D. R. LC3-associated phagocytosis at a glance. Journal of cell science 132, doi:10.1242/jcs.222984 (2019).

31. Heckmann, B. L., Boada-Romero, E., Cunha, L. D., Magne, J. & Green, D. R. LC3-Associated Phagocytosis and Inflammation. Journal of molecular biology 429, 3561–3576, doi:10.1016/j.jmb.2017.08.012 (2017).

32. Green, D. R., Oguin, T. H. & Martinez, J. The clearance of dying cells: table for two. Cell death and differentiation 23, 915–926, doi:10.1038/cdd.2015.172 (2016).

33. Filimonenko, M. et al. The selective macroautophagic degradation of aggregated proteins requires the PI3P-binding protein Alfy. Molecular cell 38, 265–279, doi:10.1016/j.molcel.2010.04.007 (2010).

34. Fox, L. M. et al. Huntington’s Disease Pathogenesis Is Modified In Vivo by Alfy/Wdfy3 and Selective Macroautophagy. Neuron 105, 813–821.e816, doi:10.1016/j.neuron.2019.12.003 (2020).

35. Simonsen, A. et al. Alfy, a novel FYVE-domain-containing protein associated with protein granules and autophagic membranes. Journal of cell science 117, 4239–4251, doi:10.1242/jcs.01287 (2004).

36. Lystad, A. H. et al. Structural determinants in GABARAP required for the selective binding and recruitment of ALFY to LC3B-positive structures. EMBO reports 15, 557–565, doi:10.1002/embr.201338003 (2014).

37. Hanada, T. et al. The Atg12-Atg5 conjugate has a novel E3-like activity for protein lipidation in autophagy. The Journal of biological chemistry 282, 37298–37302, doi:10.1074/jbc.C700195200 (2007).

38. Fujita, N. et al. An Atg4B mutant hampers the lipidation of LC3 paralogues and causes defects in autophagosome closure. Molecular biology of the cell 19, 4651–4659, doi:10.1091/mbc.E08-03-0312 (2008).

39. Martinez, J. et al. Microtubule-associated protein 1 light chain 3 alpha (LC3)-associated phagocytosis is required for the efficient clearance of dead cells. Proceedings of the National Academy of Sciences of the United States of America 108, 17396–17401, doi:10.1073/pnas.1113421108 (2011).

40. Martinez, J. Detection of LC3-Associated Phagocytosis (LAP). Curr Protoc Cell Biol 87, e104, doi:10.1002/cpcb.104 (2020).

41. Eenjes, E., Dragich, J. M., Kampinga, H. H. & Yamamoto, A. Distinguishing aggregate formation and aggregate clearance using cell-based assays. Journal of cell science 129, 1260–1270, doi:10.1242/jcs.179978 (2016).

42. Fan, J. et al. ASEP: Gene-based detection of allele-specific expression across individuals in a population by RNA sequencing. PLoS genetics 16, e1008786, doi:10.1371/journal.pgen.1008786 (2020).

43. Takahashi, Y. et al. Bif-1 interacts with Beclin 1 through UVRAG and regulates autophagy and tumorigenesis. Nature cell biology 9, 1142–1151, doi:10.1038/ncb1634 (2007).

44. Chandra, M. et al. Classification of the human phox homology (PX) domains based on their phosphoinositide binding specificities. Nature communications 10, 1528, doi:10.1038/s41467-019-09355-y (2019).

45. Wartosch, L., Gunesdogan, U., Graham, S. C. & Luzio, J. P. Recruitment of VPS33A to HOPS by VPS16 Is Required for Lysosome Fusion with Endosomes and Autophagosomes. Traffic (Copenhagen, Denmark) 16, 727–742, doi:10.1111/tra.12283 (2015).

46. Doench, J. G. Am I ready for CRISPR? A user’s guide to genetic screens. Nature reviews. Genetics 19, 67–80, doi:10.1038/nrg.2017.97 (2018).

47. Moon, H. et al. Crbn modulates calcium influx by regulating Orai1 during efferocytosis. Nature communications 11, 5489, doi:10.1038/s41467-020-19272-0 (2020).

48. Zhang, H. et al. Functional analysis and transcriptomic profiling of iPSC-derived macrophages and their application in modeling Mendelian disease. Circulation research 117, 17–28, doi:10.1161/circresaha.117.305860 (2015).

49. Chang, C. F. et al. Erythrocyte efferocytosis modulates macrophages towards recovery after intracerebral hemorrhage. The Journal of clinical investigation 128, 607–624, doi:10.1172/jci95612 (2018).

50. Wang, Y. et al. Mitochondrial Fission Promotes the Continued Clearance of Apoptotic Cells by Macrophages. Cell, doi:10.1016/j.cell.2017.08.041 (2017).

51. Patro, R., Duggal, G., Love, M. I., Irizarry, R. A. & Kingsford, C. Salmon provides fast and bias-aware quantification of transcript expression. Nature methods 14, 417–419, doi:10.1038/nmeth.4197 (2017).

52. Soneson, C., Love, M. I. & Robinson, M. D. Differential analyses for RNA-seq: transcript-level estimates improve gene-level inferences. F1000Res 4, 1521, doi:10.12688/f1000research.7563.2 (2015).

53. Love, M. I., Huber, W. & Anders, S. Moderated estimation of fold change and dispersion for RNA-seq data with DESeq2. Genome biology 15, 550, doi:10.1186/s13059-014-0550-8 (2014).

54. Zhu, A., Ibrahim, J. G. & Love, M. I. Heavy-tailed prior distributions for sequence count data: removing the noise and preserving large differences. Bioinformatics (Oxford, England) 35, 2084–2092, doi:10.1093/bioinformatics/bty895 (2019).

55. Reimand, J. et al. Pathway enrichment analysis and visualization of omics data using g:Profiler, GSEA, Cytoscape and EnrichmentMap. Nature protocols 14, 482–517, doi:10.1038/s41596-018-0103-9 (2019).

56. Subramanian, A. et al. Gene set enrichment analysis: a knowledge-based approach for interpreting genome-wide expression profiles. Proceedings of the National Academy of Sciences of the United States of America 102, 15545–15550, doi:10.1073/pnas.0506580102 (2005).

