## Supplementary Information for "A Genome-wide CRISPR Screen Identifies WDFY3 as a Novel Regulator of Macrophage Efferocytosis"

#### Summary

#### List of Supplementary Tables

**Supplementary Table 1:**
MAGeCK Analysis of CRISPR Screening of Macrophage Efferocytosis: Input vs. Non-eaters

**Supplementary Table 2:**
MAGeCK Analysis of CRISPR Screening of Macrophage Efferocytosis: Input vs. Efficient Eaters

**Supplementary Table 3:**
MAGeCK Analysis of CRISPR Screening of Macrophage Efferocytosis: non-eaters vs. efficient eaters

**Supplementary Table 4**:
Canonical Pathway Analysis of Positive Regulators by Ingenuity Pathway Analysis (IPA)

**Supplementary Table 5**:
Canonical Pathway Analysis of Negative Regulators by IPA

**Supplementary Table 6:**
Comparing the Top-ranked Positive Regulators by Our Screen and Previous Screens in U937 Monocytic Cell Line-derived Macrophages

**Supplementary Table 7:**
DESeq2 Results of RNA-seq of $Cre^-$ and $Cre^+$ BMDMs

**Supplementary Table 8**:
GSEA Output of Human Reactome Pathways Enriched in Upregulated Genes in $Cre^+$ BMDMs

**Supplementary Table 9**:
GSEA Output of Gene Ontology Biological Process Terms Enriched in Upregulated Genes in $Cre^+$ BMDMs

**Supplementary Table 10**:
GSEA Output of Human Reactome Pathways Enriched in Downregulated Genes in $Cre^+$ BMDMs

**Supplementary Table 11**:
GSEA Output of Gene Ontology Biological Process Terms Enriched in Downregulated Genes in $Cre^+$ BMDMs

#### List of Supplementary Videos

##### Supplementary Video 1: Synchronized F-actin polymerization and depolymerization during efferocytosis.

Bone-marrow-derived macrophages (BMDMs) from wild-type C57BL/6J mice were transfected with LifeAct-tagGFP2 mRNA encoding a fusion protein of LifeAct and a GFP reporter for visualization of cellular F-actin. The transfected BMDMs were then stained with CellTracker Deep Red, a fluorescent dye that freely passes through cell membranes and is well-retained in cells, allowing labeling of cytoplasmic area and tracking cell movements. BMDMs undergoing efferocytosis of unlabeled apoptotic Jurkat cells were imaged with a Nikon Ti Eclipse inverted microscope for spinning-disk confocal microscopy equipped with a 60x/1.49 Apo TIRF oil immersion lens. Time-lapse images of the same fields were acquired at 30 s intervals for 20 min, and the movie was prepared by FIJI. As pointed by the white arrow, the phagocytic cup formation is visualized by emerged F-actin signals, followed by engulfment of the apoptotic cell initially surrounded by F-actin. The complete engulfment is synchronized by reduced F-actin signals, indicating depolymerization.

### Supplementary Figures and Figure Legends

#### Supplementary Fig. 1

a

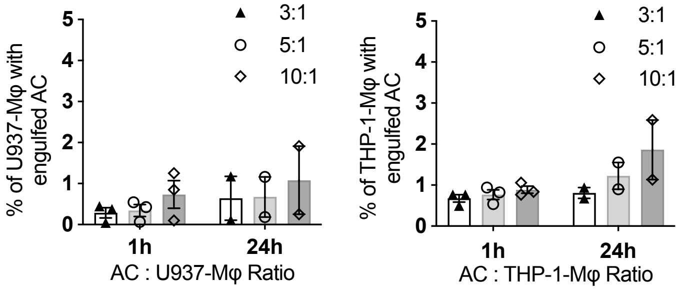

b

| | Day 0 BM Cells | Day 6 After Selection | Transduction Rate (Day 6/Day 0 x 100%) | Day 9 BMDM | Before Efferocytosis | Sorted Non-Eaters | Sorted Good Eaters |
|---|---|---|---|---|---|---|---|
| Replicate 1 | 400 x 10^6 | 138 x 10^6 | 34.5% | 140 x 10^6 | 40 x 10^6 | 15.0 x 10^6 | 2.5 x 10^6 |
| Replicate 2 | 500 x 10^6 | 126 x 10^6 | 25.2% | 130 x 10^6 | 34 x 10^6 | 16.6 x 10^6 | 3.5 x 10^6 |

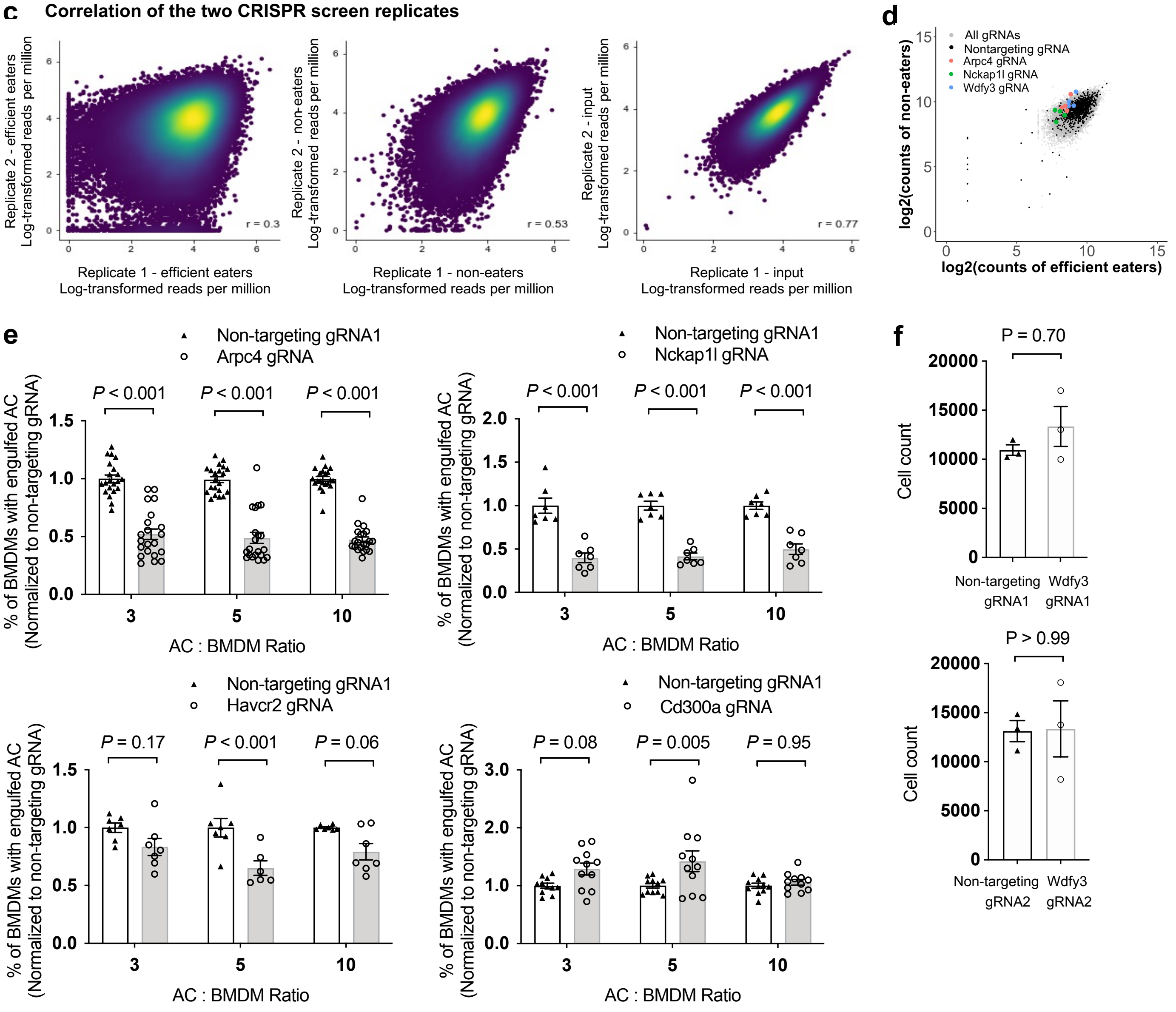

**Supplementary Fig. 1 Quality control of our genome-wide CRISPR screen and validation of selected top hits.**

**(a)** Efferocytosis of PKH26-labeled human Jurkat apoptotic cells by U937-derived macrophages and THP-1-derived macrophages (n = 2 or 3 individual experiments). **(b)** The number of cells in each step for each of the two replicates of CRISPR screens. **(c)** Pearson's correlation coefficient for gRNA counts between the two replicates for "efficient eaters", "non-eaters", and the "input" samples. **(d)** Visualization of gRNA counts of all gRNAs, the non-targeting gRNAs, and the gRNAs targeting the top positive regulators. **(e)** High Content Imaging-based individual validation of top screen hits using gRNAs in the original screening library (n = 7 independent experiments for *Arpc4*, n = 3 for *Cd300a*, n = 2 for *Nckap1l* and *Havcr2*, each from the average of 3 or 4 technical replicates). **(f)** Knockout by gRNAs targeting *Wdfy3* did not affect BMDM viability.

### Supplementary Fig. 2

| | | | | | | | | | | |
|---|---|---|---|---|---|---|---|---|---|---|
| **Source:** | This study | Haney et al., 2018 | | | | | | | | Kamber et al., 2021 |
| **Analysis Package:** | MAGeCK | casTLE | | | | | | | | casTLE |
| **Sorting method:** | FACS-based sorting | Magnetic beads-based sorting | | | | | | | | FACS-based sorting |
| **Screen library:** | Brie Library | Customed genome-wide CRISPR/Cas9 deletion library (Morgens et al., 2017) | | | | | | | | Customed genome-wide CRISPR/Cas9 deletion library (Morgens et al., 2017) |
| **Phagocyte:** | BMDMs | U937-derived macrophages | | | | | | | | J774 macrophages |
| **Substrates:** | Apoptotic Jurkat cells (~ 10 µm) | IgG-opsonized RBCs (~ 4 µm) | C3b-opsonized RBCs (~ 4 µm) | Beads (4 µm, negatively charged) | Beads (1.3 µm, negatively charged) | Beads (0.3 µm, negatively charged) | Beads (1.3 µm, positively charged) | Zymosan (~ 1 µm) | Myelin (500 nm) | APMAP$^{KO}$ Ramos cells (~ 10 µm) |
| **Rank** | | | | | | | | | | |
| 1 | U2AF1 | PRKCD | NCKAP1L | NHLRC2 | NHLRC2 | NHLRC2 | NHLRC2 | MAPK1 | MAPK1 | FERMT3 |
| 2 | ARPC4 | NCKAP1L | PRKCD | TM2D2 | NCKAP1L | TM2D2 | ACTR2 | PRKCD | ITGB2 | ITGB2 |
| 3 | DYRK1A | PRKCB | MAPK1 | LCMT1 | ABI1 | TM2D3 | NCKAP1L | NHLRC2 | NHLRC2 | TLN1 |
| 4 | WDR62 | ABI1 | ABI1 | SYS1-DBNDD2 | DOCK2 | NCKAP1L | ACTR3 | TLN1 | TLN1 | NCKAP1L |
| 5 | POMT2 | MAPK1 | DOCK2 | TLE3 | CD93 | ARPC4-TTLL3 | ABI1 | PLEK | PRKD2 | HSP90B1 |
| 6 | PREX2 | PRKD2 | BRK1 | TM2D3 | CYFIP1 | LCMT1 | DOCK2 | PRKD2 | PRKCB | WASF2 |
| 7 | HAVCR2 | DOCK2 | CREBBP | ABI1 | LCMT1 | TM2D1 | CYFIP1 | ITGB2 | PRKCD | OTUD5 |
| 8 | BCL2L13 | AP002348.1 | FBXO11 | PTPN7 | ACTR2 | PPME1 | ARPC2 | FERMT3 | SHOC2 | BRK1 |
| 9 | TRAF3 | IKBIP | PRKCB | RPS6KA1 | BRK1 | RPS6KA1 | ARPC4-TTLL3 | PRKCB | FERMT3 | GPR84 |
| 10 | WDFY3 | TCP1 | CYFIP1 | STUB1 | STUB1 | CHIC2 | ARPC4 | CERS2 | SUCNR1 | ITGAM |
| 11 | VPS52 | SGMS1 | SHOC2 | NCKAP1L | ARPC2 | KAT6A | SPPL3 | NCKAP1L | PLEK | CNOT2 |
| 12 | SEC61A1 | MTDH | NPRL2 | KAT6A | RP11-45M22.4 | HSD17B12 | LCMT1 | SHOC2 | RP11-45M22.4 | ARPC4 |
| 13 | KPNA6 | MOBP | JUNB | OTUB1 | TM2D2 | DDA1 | RAC1 | SUCNR1 | NCKAP1L | MESDC2 |
| 14 | NCKAP1L | BRK1 | AHSA1 | TBL1XR1 | PPME1 | ABI1 | ARPC3 | ABI1 | ARAF | RAB5C |
| 15 | FCRL1 | FBXW12 | PIK3R5 | PPME1 | WASF2 | CYFIP1 | MAPK1 | JUNB | MEF2D | ARPC1B |
| 16 | VARS1 | ZNF286B | AMBRA1 | ARFRP1 | LAMTOR4 | HSPA13 | AMBRA1 | FADD | ABI1 | UBR5 |
| 17 | CEBPB | TMEM256-PLSCR3 | DEPDC5 | SYS1 | ARPC3 | STUB1 | MYO9B | ELOVL1 | JUNB | COMMD2 |
| 18 | TMEM200A | CREBBP | SGMS1 | UBE2D3 | ACTR3 | CUL3 | RPL21 | RIT1 | ARPC4-TTLL3 | MEMO1 |
| 19 | ACTR3 | NPRL2 | ELOVL1 | JMJD6 | ARF1 | UBE2D3 | AIP | RP11-45M22.4 | MESDC2 | ACTR2 |
| 20 | SLC2A12 | FIS1 | TLN1 | XPR1 | PPP2R1A | CD93 | FADD | DOCK2 | C16ORF72 | DEK |
| 21 | TSC22D2 | PAOX | RAC1 | ELAVL3 | TM2D3 | AFF2 | JAK1 | BRAP | ERP44 | MAPK14 |
| 22 | ABHD17A | ARPC4 | NPRL3 | USE1 | ARPC4-TTLL3 | ACTR3 | WASF2 | GAB1 | ARPC3 | ABI1 |
| 23 | RAC1 | NARG2 | PRKD2 | C1ORF43 | LAMTOR2 | IQCA1 | RRAGA | ACTR3 | RIT1 | RABGGTA |
| 24 | PREX1 | FAM27D1 | SUCNR1 | CYFIP1 | WDR81 | TLE3 | KLF6 | FBXO11 | HSP90B1 | EED |
| 25 | SH3GLB1 | JUN | SRM | MTA2 | SASH3 | BRD9 | TM2D1 | STT3A | KRAS | ARHGAP30 |
| 26 | SNX24 | AC022498.1 | E2F8 | GAPDH | FNIP2 | GFM1 | BIN2 | TRAPPC2L | JAK1 | TMEM189 |
| 27 | CCDC115 | TIMM17A | MORF4L2 | WAS | TM2D1 | MLLT1 | BRK1 | ACTR5 | AHSA1 | FRYL |
| 28 | SETD1B | LENEP | CERS2 | DOT1L | ACTB | SLC25A51 | LAMTOR4 | ARNT | CERS2 | SMARCB1 |
| 29 | ABCE1 | YOD1 | RIT1 | CHIC2 | SAP130 | TMEM167A | RP11-45M22.4 | JAK1 | ELOVL1 | UBAP2L |
| 30 | CYFIP1 | ERP44 | EZH1 | MCM3 | RRAGA | CAND1 | TM2D2 | CSF2RB | RC3H1 | TFAP2A |
| 31 | PIAS1 | CYFIP1 | ERP44 | PCNX | RAC1 | UBE2L3 | BRD2 | ARPC2 | RAF1 | 9130011E15RIK |
| 32 | DCLRE1A | ZNF22 | ARPC4-TTLL3 | TM2D1 | PPP6R1 | RPL28 | XPR1 | ERP44 | GNAI2 | CITED2 |
| 33 | GIMAP4 | BASP1 | PLEK | C11ORF73 | ARPC4 | C11ORF73 | PTDSS1 | ACTR2 | RRAGA | GM20431 |
| 34 | CEBPA | YPEL5 | SASH3 | PTPRC | TMEM208 | RFC3 | LAMTOR2 | CREBBP | ACTR2 | BHLHE41 |
| 35 | RTL8A | LOR | RPS6KA3 | ANAPC2 | SPTLC1 | ARPC2 | PIK3R5 | AMBRA1 | GAB1 | ITGB1 |
| 36 | PRAMEF6 | MAP3K4 | DYRK1A | CD93 | LAMTOR3 | CCM2 | OTUD5 | AHR | ACTR3 | KDM6A |
| 37 | TAS2R16 | TBC1D29 | C16ORF72 | FURIN | CERS2 | NSMCE1 | C1GALT1C1 | BSG | ARPC4 | NCSTN |
| 38 | ATP1B3 | ECHDC3 | SAMD7 | MYO9B | JKAMP | ZMYND8 | UBE2D3 | LRRC8A | AMBRA1 | NRAS |
| 39 | ST8SIA2 | RBM4B | ARAF | LDLR | MYO9B | ARPC3 | HRC | ARAF | UBLCP1 | FBXW7 |
| 40 | COMMD4 | RAX2 | WASF2 | BCL11A | PCNX | NPEPPS | USP34 | SPPL3 | RPS6KA3 | SOS1 |
| 41 | SPTY2D1 | ELOVL1 | YPEL5 | KCTD5 | TIMMDC1 | LBX1 | GNAI2 | PPP1CC | STT3A | CCM2 |
| 42 | WASF2 | TRIM41 | WDR3 | PPP6R1 | NARS2 | WDR36 | BASP1 | CYFIP1 | RRAGC | METTL16 |
| 43 | ABI1 | AHSA1 | STT3B | LZTR1 | SPTSSA | LZTR1 | WDR81 | RRAGA | CSK | UCHL5 |
| 44 | TREM2 | ELOVL5 | SETD1B | GFI1 | BASP1 | CDC42 | ATAD2 | ARPC3 | MAP2K3 | ST3GAL3 |
| 45 | SHLD1 | TRNP1 | KDM1B | CEBPE | HOXD13 | KAT7 | SF3B1 | KRAS | ACTB | ZMYM4 |
| 46 | GTF3C1 | GADD45B | LRRC8A | TBC1D10B | ARF6 | FLI1 | ZCCHC7 | TRBV27 | CREBBP | FBXO11 |
| 47 | TLE4 | GSTT2B | MDN1 | VPS54 | CAPN9 | C15ORF53 | PACS2 | RPS6KA3 | NIFK | SLC25A26 |
| 48 | ISY1 | MEP1A | SNRNP25 | NF1 | COX20 | PCNX | HECTD1 | SLC35B4 | BRAP | RAB1 |
| 49 | OR10K1 | AL591684.1 | HNRNPF | TMEM203 | NGLY1 | PPP2R2A | PDCL | DAZAP1 | ACTR5 | UGP2 |
| 50 | CDK9 | MRPL2 | SND1 | SPRED2 | UBE2A | CHD2 | BCL6 | C19ORF25 | LAMTOR2 | ARPC2 |
| 51 | MGAT1 | CDKL1 | RAF1 | RAB10 | MESDC2 | PTPN7 | MRPL18 | QKI | RAB7A | YTHDF3 |
| 52 | VPS33A | FKBP5 | THRAP3 | KIAA1109 | UBE2D3 | CAB39 | TOR1B | TRAF6 | OSTC | PSMA1 |
| 53 | ACTR2 | TRIM48 | ARL4A | SLC38A2 | CCER1 | ANLN | C5ORF47 | C16ORF72 | DOCK2 | KRAS |
| 54 | LACTB | FEZF2 | PRKRA | PAXBP1 | CHMP5 | STK11 | CDK11A | MEF2D | CYFIP1 | YWHAE |
| 55 | FBXW12 | FBXO11 | KLF16 | RPL28 | C1GALT1C1 | ARF4 | TMEM30A | GNAI2 | ZNF699 | CYFIP1 |
| 56 | ELOVL5 | RAC1 | ITFG2 | ABT1 | NFE2L2 | TBCB | VPS4B | IRF8 | ELF1 | MLLT1 |
| 57 | HNRNPK | ZNF699 | ATAD1 | PRELID1 | LSMD1 | AKT2 | C1ORF220 | APEH | BRK1 | INTS10 |
| 58 | SF3B2 | SARNP | CTSE | NAPG | MAFA | UBE2K | EWSR1 | MAP2K3 | ARPC2 | RUNX1 |
| 59 | NFKBIA | FJX1 | PAPD5 | PRPH2 | AC021860.1 | RSRC2 | SFTPA1 | LARP4 | YPEL5 | SPOP |
| 60 | FBXW2 | TIMM13 | BIN2 | SBDS | MAB21L3 | AAGAB | ACAD8 | AIP | SPCS2 | ZKSCAN5 |
| 61 | DENND3 | RNF187 | EHD3 | AFF2 | KIAA1432 | TNFAIP3 | TRAPPC2L | STIP1 | MAGT1 | SH3BP2 |
| 62 | TMEM220 | CKAP4 | CARM1 | BRK1 | GPS2 | ANAPC10 | RAX | ARPC4-TTLL3 | KSR1 | WAS |
| 63 | CCDC9B | RAD51AP1 | FERMT3 | MYEOV2 | MYBPC2 | KCTD5 | ZFAND4 | CSK | BCL6 | ELAVL1 |
| 64 | ALPK1 | ALKBH2 | OR1S1 | NUDCD2 | TDRD3 | MTA2 | DNAJB4 | UBL5 | REC8 | WIPF1 |
| 65 | UROD | AKAP2 | C12ORF66 | COQ10B | RPL28 | GDF1 | ARPC5 | JAK2 | MAP2K1 | FXR1 |
| 66 | ZBTB46 | AL592284.1 | ZBTB1 | NCOR1 | FAM90A1 | WI2-3308P17.2 | RP11-146D12.2 | GPR89C | ALK | BC051142 |
| 67 | ODAD2 | PKN3 | PHF6 | SRRD | USF1 | CBWD7 | CSTF3 | CSF2RA | STT3B | PTPRO |
| 68 | SOCS5 | RARRES2 | LBX1 | NSMCE2 | TMEM60 | HIST2H2AA4 | CASP9 | PIK3R5 | SAMSN1 | LRRC25 |
| 69 | PDCD1 | AC020922.1 | STEAP3 | TUSC1 | PIK3R5 | VPS11 | WSB1 | FLCN | GABARAPL2 | OLFR1167 |
| 70 | TAS1R1 | ABHD14B | GRPEL1 | CCNH | HIST1H2BJ | RPRD2 | SASH3 | RREB1 | FOXJ3 | PNRC1 |
| 71 | FIZ1 | AK2 | FRG2C | CCNF | SERP1 | TFEB | PDHA1 | UTP15 | RAX | DSTYK |
| 72 | GRIN2D | TP53 | NXF5 | TNPO1 | AMBRA1 | C1ORF43 | KDM2A | BCL6 | CELF2 | CLASRP |
| 73 | TOPAZ1 | C2ORF27B | KPTN | NIP7 | SNRK | RAB1A | AKT1 | KDM1B | CTTNBP2 | CFP |
| 74 | ARPC3 | RP11-204N11.1 | RC3H1 | JKAMP | HIST1H2AH | WDR7 | FMO5 | MS4A8 | MSI1 | XPO6 |
| 75 | MTMR9 | SULT1C2 | SKA1 | WASF2 | NDUFA4 | TMEM203 | GOLGA8S | LAMTOR3 | GSK3B | FLCN |
| 76 | TMEM98 | NKD2 | MAF1 | ZMYND8 | ELMO2 | MYO9B | CERS2 | FNIP2 | OR2D3 | SURF2 |
| 77 | FBXO11 | PYURF | KATNA1 | LIMS3L | SERBP1 | ACTR2 | KLK14 | DEPDC5 | NDUFA4L2 | SUZ12 |
| 78 | CCDC88B | LCE1F | SPAM1 | TVP23C-CDRT4 | TFEB | AC009060.1 | EHD1 | OR4A16 | FAM83A | WDR26 |
| 79 | WNK1 | CYS1 | PTPLB | MIDN | C21ORF128 | ZFR | PPP2R2A | RAB7A | IGSF9 | TADA2B |
| 80 | CEACAM4 | HSBP1L1 | ITIH1 | GTPBP3 | TCERG1 | ZC3HC1 | PROCA1 | AHSA1 | FBXO11 | RAB14 |
| 81 | SERPINB3 | TMEM11 | ELF4 | ZC3HC1 | FERMT2 | ZFX | PKN1 | API5 | SETD1B | CRAMP1L |
| 82 | OPA3 | CDKN2C | ERG | FTO | CCNF | PA2G4 | APBB3 | AC025278.1 | WTAP | PIK3CB |
| 83 | CTR9 | DUS3L | ARGLU1 | PDHA1 | STT3A | MYC | PITRM1 | BIN2 | FGL1 | MAFB |
| 84 | PRL | ZNF503 | SFPQ | KIF11 | TLN1 | COX8A | ELOVL1 | RP11-512M8.5 | WASF2 | ZBTB7A |
| 85 | U2AF2 | AL645730.2 | USP46 | SAP130 | ARFRP1 | ELAVL1 | BRAP | CBLC | LAMTOR4 | ZFP644 |
| 86 | METTL16 | GORASP1 | RNF133 | EWSR1 | HSP90B1 | CSNK2A2 | KIAA1109 | SCD | PHACTR4 | PIK3R1 |
| 87 | POT1 | PIK3R6 | TRIM48 | RAX | BRD7 | PET112 | USP17L2 | RRAGC | SPPL3 | DNTTIP1 |
| 88 | APOL4 | KCTD13 | NHLRC2 | XPO6 | NOC2L | PEPD | CSTF2 | FAM203A | CTNND2 | ADRBK1 |
| 89 | APOL1 | GJA3 | TECPR2 | AC019206.1 | LLPH | XPR1 | PPIAL4D | PTPLB | LAMTOR3 | CSE1L |
| 90 | APOL3 | C5ORF20 | BRD2 | REG1A | WAS | LAMTOR2 | SEC23B | GNB1 | ZSWIM8 | GNB2 |
| 91 | APOL2 | CYBB | DNAH5 | FBXW12 | NUDCD2 | STAM | TRAF2 | TECR | ICMT | PREX1 |
| 92 | KATNBL1 | MRPS18A | BRAP | RP11-763F8.1 | TPRKB | RASA2 | XRCC6BP1 | STT3B | FNIP2 | STRIP1 |
| 93 | TDO2 | NAA15 | EIF5B | ELAVL1 | DAXX | NUDCD2 | SRRT | FOXJ3 | BIN2 | SNX17 |
| 94 | RHO | AL359091.2 | MAPK14 | KIAA1432 | APRT | CCNF | KIDINS220 | GADD45B | CNOT11 | RELA |
| 95 | THAP3 | RAB13 | CCDC6 | NEDD8 | WAPAL | PDHB | DDX28 | TRAF3IP3 | PLA2G4E | MAP4K4 |
| 96 | NPY4R | AC112721.1 | MUL1 | WDR81 | AUP1 | CDC73 | PGAM1 | RIC8A | CDC42 | DPH2 |
| 97 | NPY4R2 | JUNB | TCEANC | AMBRA1 | DHTKD1 | ZGLP1 | MTHFD1 | ELOVL5 | ADRM1 | MATK |
| 98 | LYG2 | HIST1H1E | SZT2 | SNRK | HSA-MIR-150 | ACKR4 | CDC42 | SFPQ | BOD1 | TMEM237 |
| 99 | ATP6V0B | SUN1 | CHMP3 | UNC50 | FLCN | COX6B1 | CREBBP | ELF4 | RASGRP3 | ARPC3 |
| 100 | RPS23 | RNF38 | MESDC2 | CHMP6 | HAUS7 | HNRNPA2B1 | RPSA | MYD88 | SASH3 | OLFR389 |

**Frequency**

| Frequency |
|---|
| 1 |
| 2 |
| 3 |
| 4 |
| 5 |
| 6 |
| 7 |
| 8 |
| 9 |
| 10 |

**Supplementary Fig. 2 Comparing the top-ranked positive regulators by our screen and previous screens in U937 monocytic cell line-derived macrophages using an array of substrates.**

The heatmap visualizes the overlap of the top-ranked positive regulators in our screen of efferocytosis in primary macrophages and in previously published screens of phagocytosis of an array of substrates in U937-derived macrophages (Haney et al., 2018[1]). The matrix for generating the heatmap is in **Supplementary Table S6**. Murine gene symbols were converted to human gene symbols using gProfiler at https://biit.cs.ut.ee/gprofiler/orth.

**Supplementary Fig. 3**

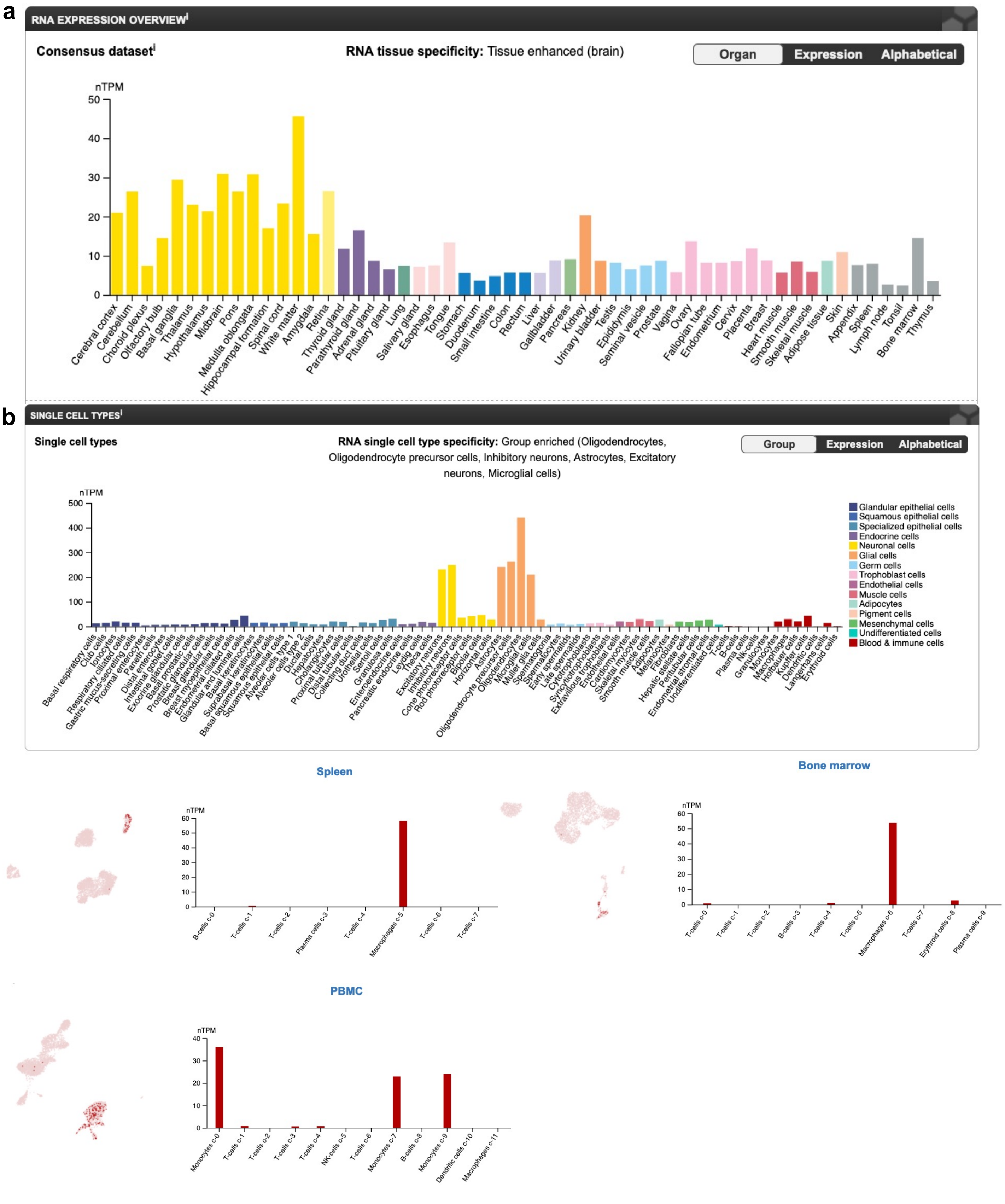

**c**

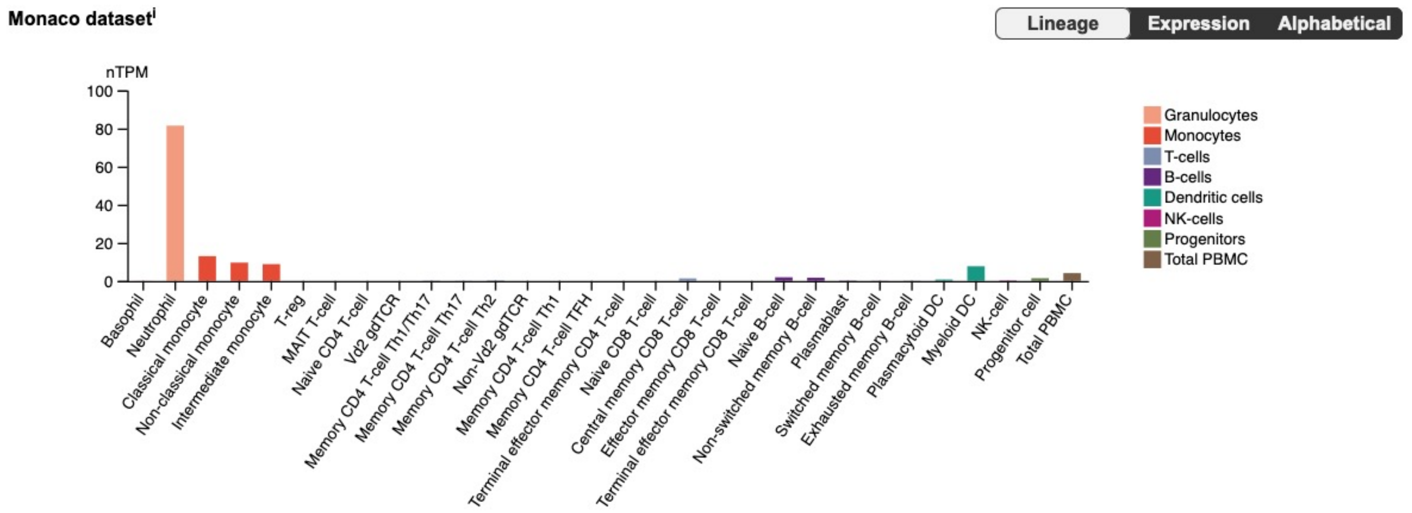

**Supplementary Fig. 3 WDFY3 expression across tissue and cell types based on data in the Human Protein Atlas resource.**

**(a)** The consensus dataset consists of normalized transcript expression (nTPM) levels for 55 tissue types, created by combining the HPA and GTEx transcriptomics datasets using the internal normalization pipeline. Color-coding is based on tissue groups, each consisting of tissues with functional features in common. **(b)** Single cell transcriptomics data for 25 tissues and peripheral blood mononuclear cells (PBMCs) were analyzed. These datasets were respectively retrieved from the Single Cell Expression Atlas, the Human Cell Atlas, the Gene Expression Omnibus, the Allen Brain Map, and the European Genome-phenome Archive. **(c)** The nTPM levels resulting from the internal normalization pipeline are visualized for 29 blood cell types and total PBMCs from Monaco et al [2]. The panels are screenshots from the Human Protein Atlas resource on November 24, 2021.

#### Supplementary Fig. 4

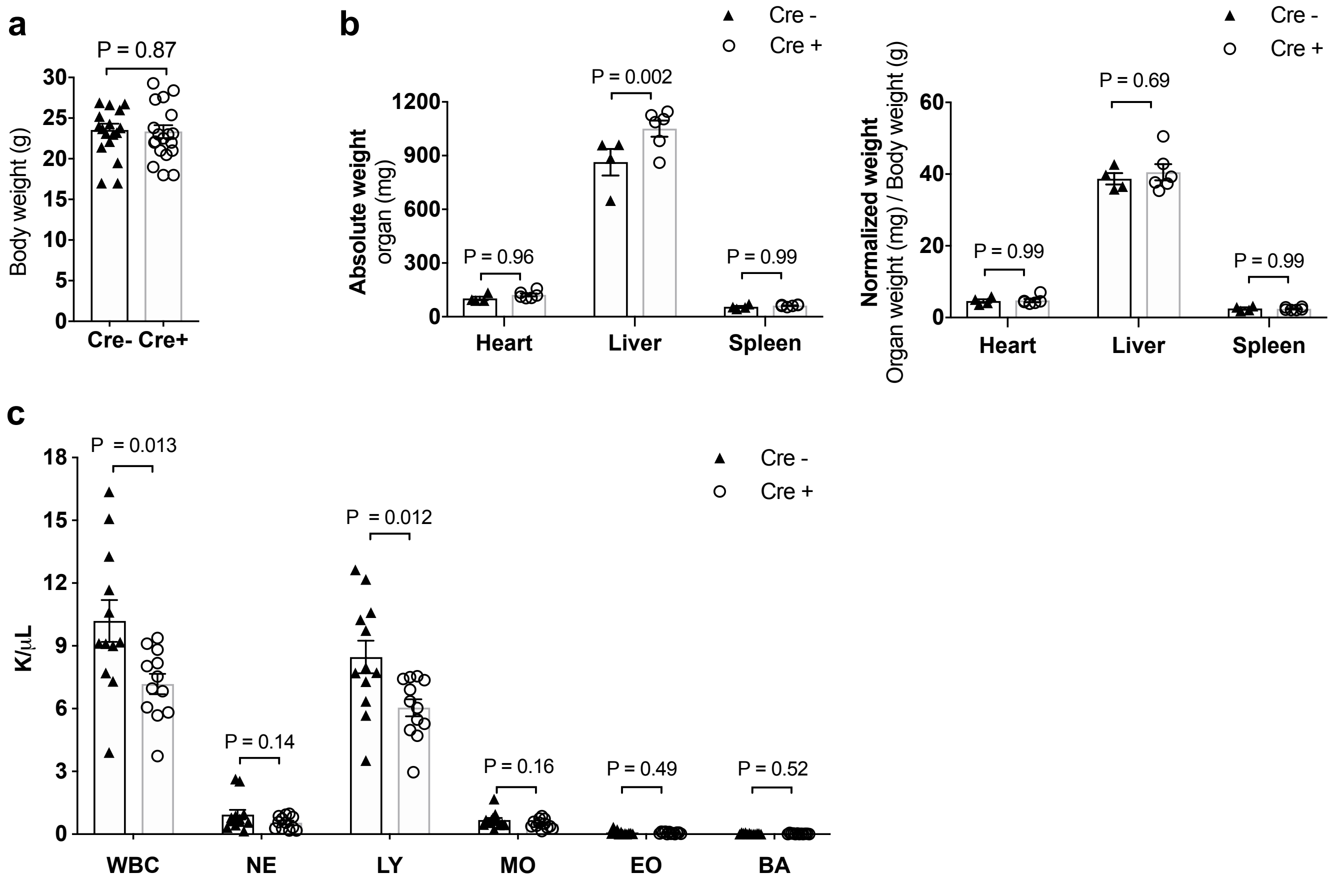

**Supplementary Fig. 4 Basic characteristics of myeloid-specific *Wdfy3* knockout mice.**

**(a)** Body weight (n = 18 or 20 biological replicates). **(b)** Organ weight and organ weight normalized to body weight (n = 6 biological replicates). **(c)** Complete Blood Count with Differential (n = 12 biological replicates).

**Supplementary Fig. 5**

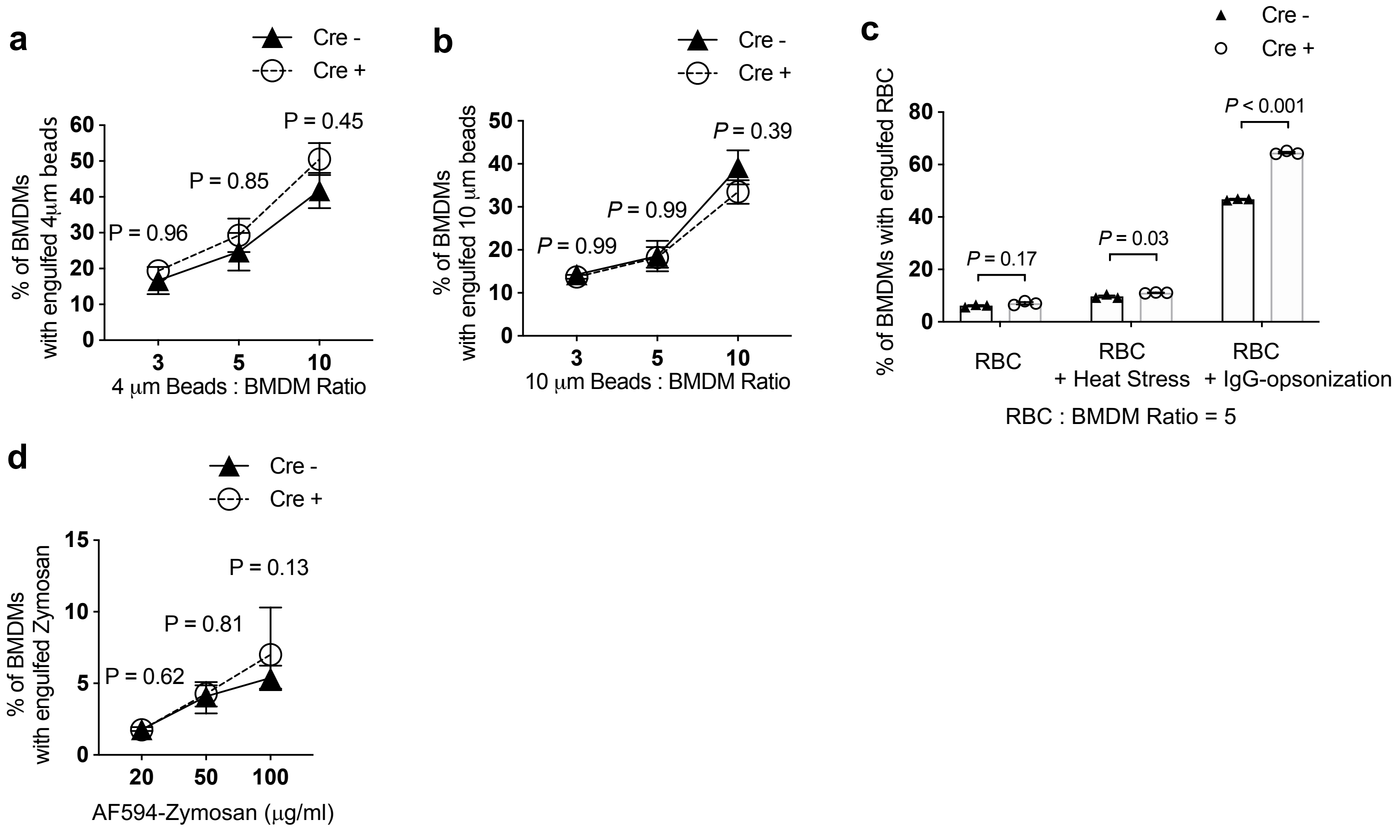

**Supplemental Fig. 5 Phagocytosis of different substrates quantified by flow cytometry.**

**(a)** 4 μm latex beads (n = 3 biological replicates with the average of 2 technical replicates). **(b)** 10 μm latex beads (n = 3 biological replicates with the average of 2 technical replicates). **(c)** Sheep red blood cells (RBCs) were either untreated, heat-stressed, or IgG-opsonized (n = 3 biological replicates with the average of 2 technical replicates). **(d)** Alexa Fluor (AF)-594 labeled Zymosan particles (n = 6 biological replicates with the average of 2 technical replicates).

**Supplementary Fig. 6**

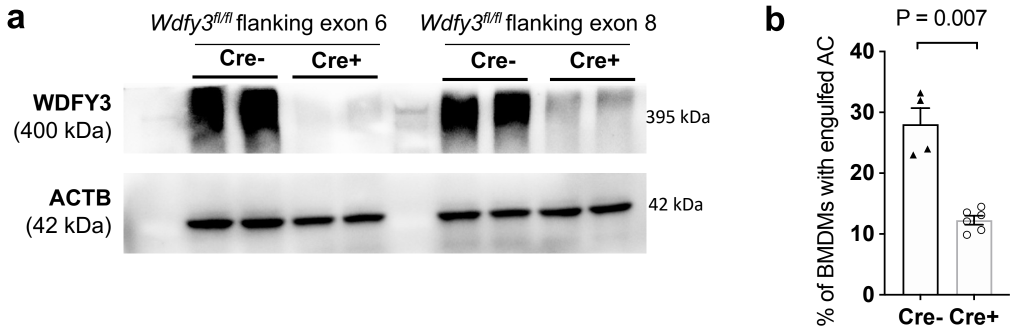

**Supplementary Fig. 6 Validation of impaired efferocytosis in BMDMs from myeloid-specific *Wdfy3* knockout mice on C57BL/6NJ background.**

**(a)** *Wdfy3*$^{fl/fl}$ mice generated by the Knock-Out Mouse Project (KOMP) with two *loxP* sites flanking exon 8, were maintained on C57BL/6N background[2]. Breeding to LysMCre mice led to efficient knockout of *WDFY3* though a small amount of residual protein remained detectable (n = 2 biological replicate). **(b)** Uptake of Hoechst-labeled apoptotic cells (ACs) was impaired in BMDMs of Cre$^{+}$ mice with Cre-lox mediated deletion of exon 8 and on C57BL/6NJ background, supporting the role *Wdfy3* in macrophage efferocytosis independent of the genetic strain and the specific gene-inactivating mutation of the mouse models (n = 4 biological replicates with the average of 2 technical replicates).

**Supplementary Fig. 7**

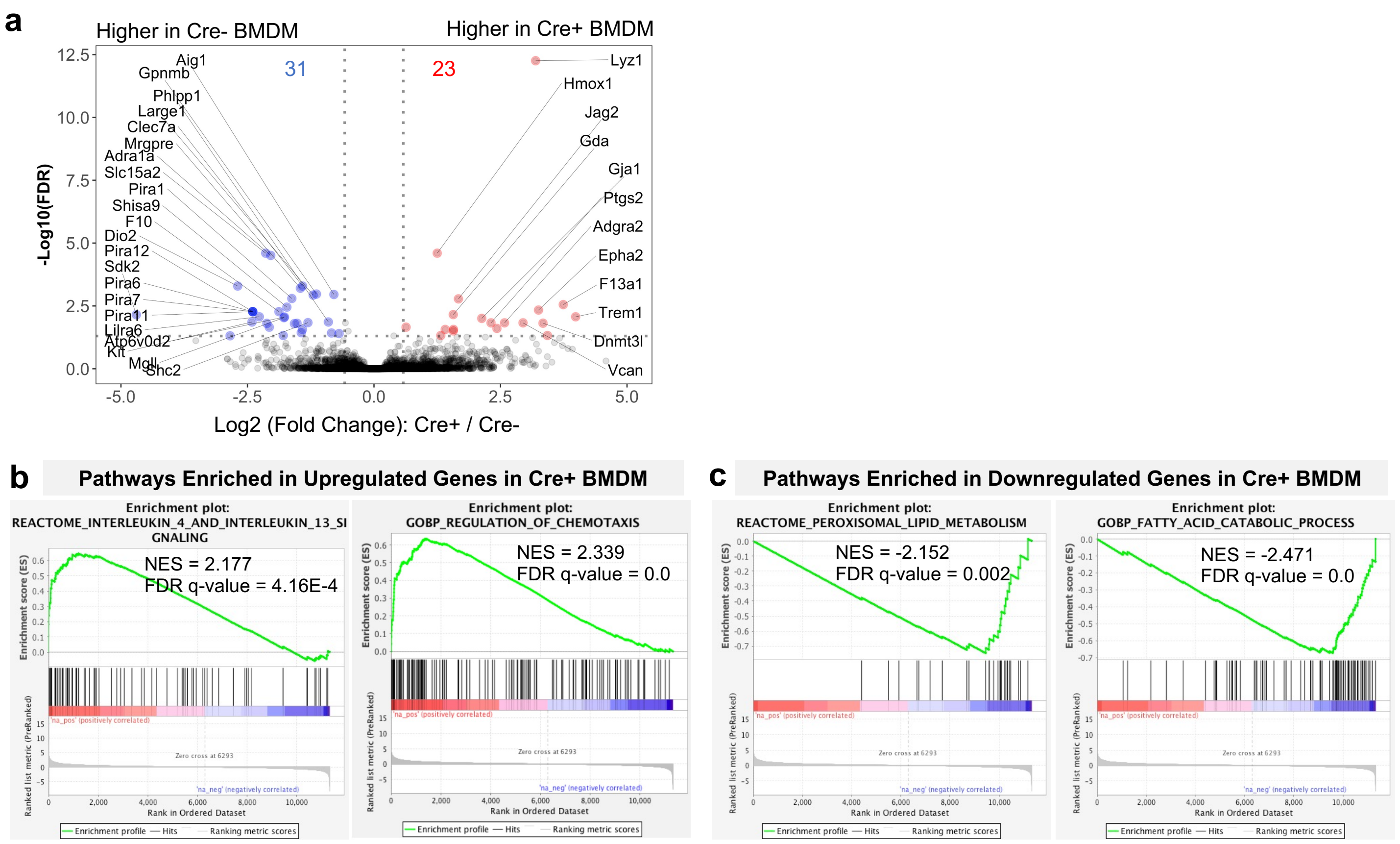

**Supplementary Fig. 7 The effects of *Wdfy3* deficiency on BMDM transcriptomic profile by RNA-seq.**

**(a)** Volcano plot highlights differentially expressed (DE) genes in Cre$^{+}$ vs. Cre$^{-}$ BMDMs (n = 4 biological replicates, male mice). **(b)** Selected top Human Reactome Pathway and Gene Ontology Biological Process terms enriched in the upregulated genes in Cre$^{+}$ BMDMs vs. Cre$^{-}$ BMDMs. Complete GSEA results are shown in **Supplementary Table 8** and **Supplementary Table 9**. **(c)** Selected top Human Reactome Pathway and Gene Ontology Biological Process terms enriched in the downregulated genes in Cre$^{+}$ BMDMs vs. Cre$^{-}$ BMDMs. Complete GSEA results are shown in **Supplementary Table 10** and **Supplementary Table 11**.

**Supplementary Fig. 8**

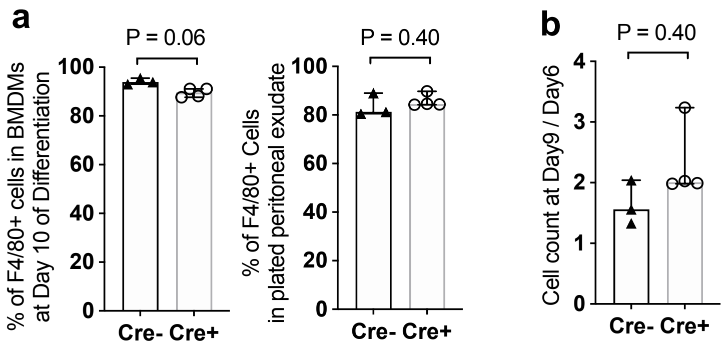

**Supplementary Fig. 8 *Wdfy3* knockout did not affect macrophage differentiation and proliferation.**

**(a)** The percentage of F4/80$^+$ macrophages in BMDMs and peritoneal exudate was comparable between Cre$^-$ and Cre$^+$ mice (n = 3 or 4 biological replicates with an average of 2 technical replicates). **(b)** Bone marrow **(**BM) cells were differentiated to BMDMs in 9 days and cell numbers were counted on day 6 and day 9. The ratio of cell counts on day 9 / day 6 implicates population doubling, indicative of proliferation capacity. The ratio shows no statistical difference between Cre$^-$ and Cre$^+$ mice (n = 3 or 4 biological replicates).

#### Supplementary Fig. 9

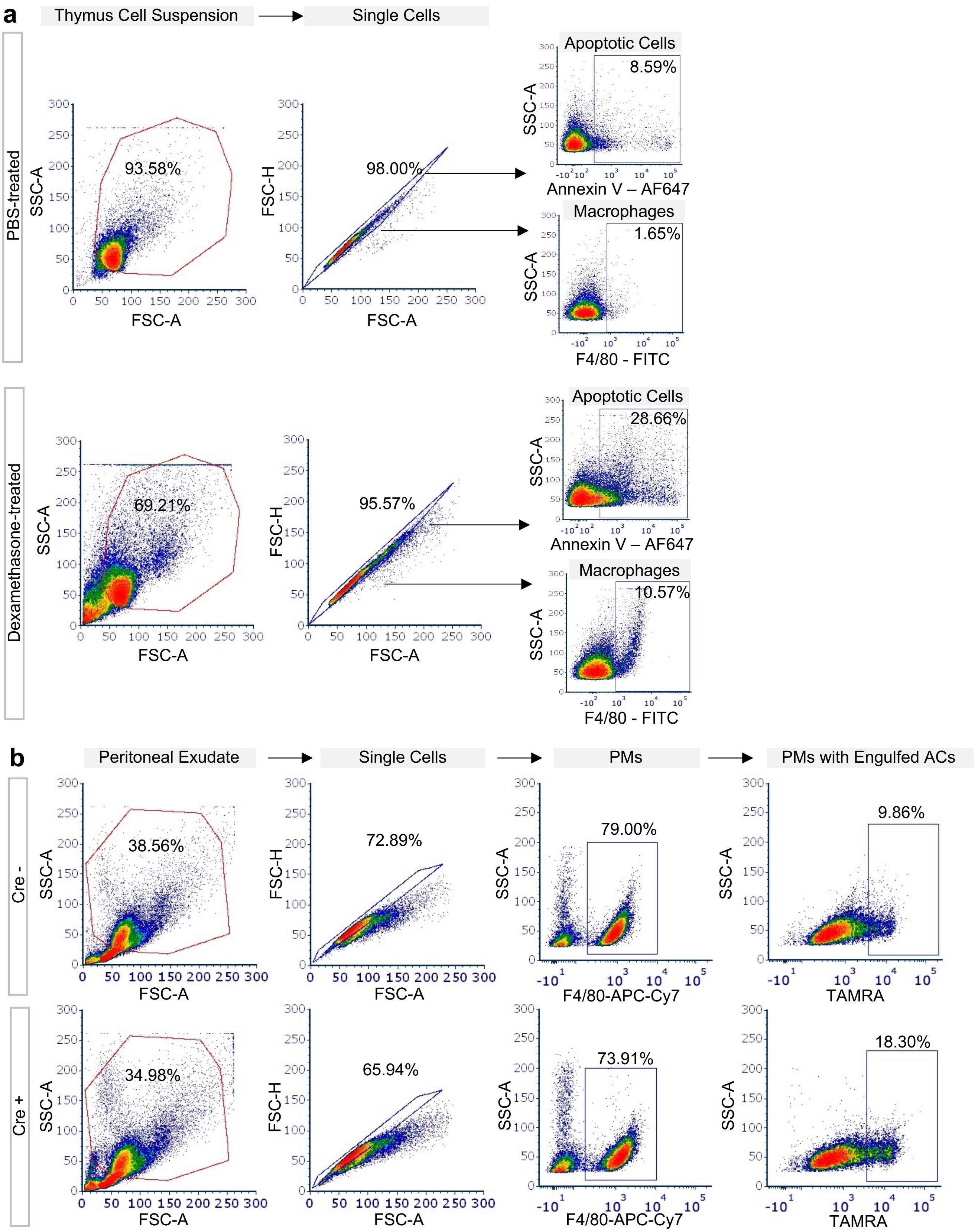

**Supplementary Fig. 9 Gating strategy of flow cytometry analysis.**

**(a)** Gating strategy of flow cytometry analysis for *in vivo* thymus efferocytosis. **(b)** Gating strategy of flow cytometry analysis for *in vivo* peritoneal macrophage efferocytosis.

#### Supplementary Methods

Cell Lines and Primary Cells:

| Reagent or Resource | Source | Identifier |
|---|---|---|
| Human: Jurkat cells | ATCC | TIB-152 |
| Human: THP-1 cells | ATCC | TIB-202 |
| Human: U937 cells | ATCC | CRL-1593.2 |
| Human: Peripheral Blood Mononuclear Cells | New York Blood Center | N/A |
| Mouse: L-929 Fibroblasts | ATCC | CCL-1 |
| Mouse: Bone Marrow-Derived Macrophages | This paper | N/A |
| Mouse: Peritoneal Macrophages | This paper | N/A |

Mice:

| Reagent or Resource | Source | Identifier |
|---|---|---|
| C57BL/6J | The Jackson Laboratory | JAX: 000664 |
| LysMCre$^{+/-}$: C57BL/6J | The Jackson Laboratory | JAX: 004781 |
| Rosa-Cas9 knockin: C57BL/6J | The Jackson Laboratory | JAX: 026179 |
| Wdfy3$^{fl/fl}$: 129/SvEv x C57BL/6 (flanking Exon 6) | Ai Yamamoto Lab<br>Dragich et al., 2016.[3] | N/A |
| Wdfy3$^{fl/fl}$: C57BL/6NJ (flanking Exon 8) | Konstantinos Zarbalis Lab<br>Orosco, L. A. et al. 2014.[4] | N/A |

#### gRNAs for CRISPR Screen Validation:

| Target | Source | gRNA sequence (5' to 3') | PAM | Target Context Sequence (5' to 3') |
|---|---|---|---|---|
| Non-targeting gRNA1 | Brie Library (Addgene Cat#73633) | GGGGTAGGCCTAATTACGGA | N/A | N/A |
| Non-targeting gRNA2 | Sigma-Aldrich (Cat# CRISPR18) | N/A | N/A | N/A |
| Wdfy3 gRNA1 | Brie Library (Addgene Cat#73633) | ATGAAGTCTGATGTCATGAG | GGG | ACCCATGAAGTCTGATGTCATGAGGGGTCG |
| Wdfy3 gRNA2 | Sigma-Aldrich | Refer to Sanger Clone ID# MM5000009433 | N/A | CATGGTGACGGAGATCCGGAGG |
| Arpc4 gRNA | Brie Library (Addgene Cat#73633) | CTTTGTAATCCACTTCATGG | AGG | TGGACTTTGTAATCCACTTCATGGAGGAGA |
| Nckap1l gRNA | Brie Library (Addgene Cat#73633) | GACACGCTGGTATATGCCTG | TGG | CTTCGACACGCTGGTATATGCCTGTGGTTG |
| Havcr2 gRNA | Brie Library (Addgene Cat#73633) | CTAAAGGGCGATCTCAACAA | AGG | CCAGCTAAAGGGCGATCTCAACAAAGGAGA |
| Cd300a gRNA | Brie Library (Addgene Cat#73633) | GGGAATAGTCATGTTCACGG | TGG | CCCTGGGAATAGTCATGTTCACGGTGGCCC |

#### Plasmids for WDFY3 Cloning:

| Reagent or Resource | Source | Identifier |
|---|---|---|
| pcDNA-myc-WDFY3$_{2543\text{-}3526}$ | Ai Yamamoto Lab Filimonenko et al., 2010.[5] | N/A |
| pLE4 | A gift from Guang-Hui Liu lab Fang et al., 2018.[6] | N/A |

Primers for Genotyping:

| Strain | Source | Forward (5' to 3') | Reverse (5' to 3') |
|---|---|---|---|
| Wdfy3$^{fl/fl}$: 129/SvEv x C57BL/6 (flanking Exon 6) | Ai Yamamoto Lab. Dragich et al., 2016.[3] | GAAAGCAAGCTCGTTTACGG | AGGTTACCAGCCACAACCAG |
| Wdfy3$^{fl/fl}$: 129/SvEv x C57BL/6 (flanking Exon 6) | Ai Yamamoto Lab. Dragich et al., 2016.[3] | ACTTGGGAAGAGGGAAGCTC | AGGTTACCAGCCACAACCAG |
| Wdfy3$^{fl/fl}$: C57BL/6NJ (flanking Exon 8) | Konstantinos Zarbalis Lab. Orosco, L. A. et al. 2014.[4] | AGTGCAAATAAAGAACTAAATTAGAAGG | CATAACTTCGTATAATGTATGCTATACG |
| Wdfy3$^{fl/fl}$: C57BL/6NJ (flanking Exon 8) | Konstantinos Zarbalis Lab. Orosco, L. A. et al. 2014.[4] | ACAGGTCTCTTTGGCTGAGG | AATGTCTTGCGTCGGAAAAG |
| LysMCre$^{-/-}$ | The Jackson Laboratory | CCCAGAAATGCCAGATTACG | CTTGGGCTGCCAGAATTTCTC |
| LysMCre$^{+/-}$ | The Jackson Laboratory | TTACAGTCGGCCAGGCTGAC | CTTGGGCTGCCAGAATTTCTC |

Primers for quantitative RT-PCR:

| Target | Source | Forward (5' to 3') | Reverse (5' to 3') |
|---|---|---|---|
| Human: *WDFY3* | OriGene | GACAACCTCTGTCTCACTCCTG | GCAAATGGTCCATCACGCTATCC |
| Human: *ACTB* | PrimerBank | CATGTACGTTGCTATCCAGGC | CTCCTTAATGTCACGCACGAT |

siRNAs

| Reagent or Resource | Source | Identifier |
|---|---|---|
| ON-TARGET plus non-targeting pool | Dharmacon | D-001810-10-05 |
| ON-TARGET plus non-targeting siRNA #1 | Dharmacon | D-001810-01-20 |
| ON-TARGET plus Human WDFY3 (23001) siRNA - SMARTpool | Dharmacon | L-012924-01-0005 |
| ON-TARGET plus Human WDFY3 (23001) siRNA - Individual | Dharmacon | J-012924-09-0010 |

Antibodies:

| **Reagent or Resource** | **Source** | **Identifier** | **Dilution and the Working Concentration** |
|---|---|---|---|
| Rabbit Anti-LC3B | Abcam | Cat# ab48394 | 1:1000, 1 µg/mL |
| Rabbit anti-LC3A/B (AF488) | Cell Signaling Technology | Cat# 13082S,<br>Lot 7 (60 ug/mL) | 1:50, 1.2 ug/mL |
| Rabbit anti-GABARAP (N-term) | Abgent | Cat# AP1821a | 1:1000 (Western), 0.025 µg/mL<br>1:100 (IP), 2.5 µg/mL |
| Rabbit anti-GABARAP | Cell Signaling Technology | Cat# 13733S,<br>Lot 4 (400 ug/mL) | 1:100 (IP), 4 ug/mL |
| Rabbit anti-WDFY3 | Ai Yamamoto Lab<br>Fox et al., 2020.[7] | N/A | 1:1000 |
| Rat anti-mouse F4/80 (FITC) | BioLegend | Cat# 123108 | 1:200, 2.5 µg/mL |
| Rat anti-mouse F4/80 (APC-Cy7) | BioLegend | Cat# 123118 | 1:200, 1 µg/mL |
| Rat anti-mouse CD68 | Abcam | Cat# ab53444 | 1:200, 5 µg/mL |
| Rabbit β-Actin (HRP) | Cell Signaling Technology | Cat# 5125S,<br>Lot 6 (48 µg/mL) | 1:5000, 0.0096 µg/mL |
| Goat anti-rabbit IgG (HRP) | EMD Millipore | Cat# AP156P | 1:5000, 0.16 µg/mL |

Cell Culture Medium:

| **Reagent or Resource** | **Source** | **Identifier** |
|---|---|---|
| Dulbecco’s Modified Eagle Media (DMEM) | Corning | Cat# 10-017-CM |
| Dulbecco's Phosphate-Buffered Salt Solution 1X (DPBS) | Corning | Cat# 21-031-CM |
| Opti-MEM I Reduced Serum Medium | Gibco | Cat# 31985070 |
| Roswell Park Memorial Institute (RPMI) 1640 Media | Corning | Cat# 10-041-CM |

| | | |
|---|---|---|
| Heat-Inactivated Fetal Bovine Serum | Gibco | Cat# 10082147 |
| CellStripper | Corning | Cat# 25-056-CI |

Chemicals and Recombinant Cytokines:

| Reagent or Resource | Source | Identifier |
|---|---|---|
| Cytochalasin D | Sigma-Aldrich | Cat# C8273 |
| Dexamethasone | Sigma-Aldrich | Cat# 265005-100MG |
| Digitonin | Sigma-Aldrich | Cat# D141-500MG |
| Human Macrophage Colony Stimulating Factor (M-CSF) | Goldbio | Cat# 1120-09-100 |
| Puromycin | Sigma-Aldrich | Cat# 540411-25MG |
| Staurosporine | Alfa Aesar | Cat# J62837-M^ |

Commercial Assay Kits:

| Reagent or Resource | Source | Identifier |
|---|---|---|
| In Situ Cell Death Detection Kit, TMR red (TUNEL) | Roche | Cat# 12156792910 |
| BCA Protein Assay Kit | Thermo Scientific | Cat# 23227 |
| High Capacity cDNA Reverse Transcription Kit | Applied Biosystems | Cat# 4368814 |
| West Pico PLUS Chemiluminescent Substrate | Thermo Scientific | 34580 |
| Quick-RNA Mini Kit | Zymo | R1055 |
| DNeasy Blood and Tissue Kit | Qiagen | 69504 |

Reagents for Efferocytosis and Phagocytosis Assays:

| Reagent or Resource | Source | Identifier |
|---|---|---|
| Annexin V Conjugates for Apoptosis Detection | Invitrogen | Cat# A23204 |
| CellTracker Green CMFDA Dye | Invitrogen | Cat# C2925 |
| CellTracker Deep Red Dye | Invitrogen | Cat# C34565 |
| Cell-Vue Claret Fluorescent Cell Linker | Sigma-Aldrich | Cat# MINCLARET-1KT |
| Diluent C | Sigma-Aldrich | Cat# CGLDIL-6X10ML |
| FluoSpheres Sulfate Microspheres, 4.0 µm, red fluorescent (580/605) | Invitrogen | F8858 |
| FluoSpheres Polystyrene Microspheres, 10 µm, orange fluorescent (540/560) | Invitrogen | F8833 |
| Hoechst 33342 Solution | Thermo Scientific | Cat# 62249 |
| HCS NuclearMask™ Blue Stain | Invitrogen | Cat# H10325 |
| PKH26 Red Fluorescent Cell Linker Kit for General Cell Membrane Labeling | Sigma-Aldrich | Cat# PKH26GL-1KT |
| PKH67 Green Fluorescent Cell Linker Kit for General Cell Membrane Labeling | Sigma-Aldrich | Cat# PKH67GL-1KT |
| pHrodo Red, succinimidyl ester | Invitrogen | Cat# P36600 |
| pHrodo Red Zymosan Bioparticles Conjugate for Phagocytosis | Invitrogen | Cat# P35364 |
| Sheep Red Blood Cells 10% washed pooled cells | Rockland Immunochemicals | R405-0050 |
| siR-actin | Cytoskeleton | Cat# CY-SC001 |
| TAMRA | Invitrogen | Cat# C1171 |
| Zymosan A (S. cerevisiae) BioParticles, Alexa Fluor 594 conjugate | Invitrogen | Cat# Z23374 |

Other Reagents and Supplies:

| Reagent or Resource | Source | Identifier |
|---|---|---|
| Ficoll-Paque Premium | Sigma-Aldrich | GE17-5442-02 |
| Pierce 16% Formaldehyde (w/v), Methanol-free | Thermo Scientific | 28908 |
| Fugene 6 | Promega | Cat# E2691 |
| Immun-Blot PVDF Membrane, 0.2 µm | Bio-rad | Cat# 1620177 |
| Lenti-X Concentrator | Clontech | Cat# 631231 |
| Lipofectamine RNAiMAX Transfection Reagent | Invitrogen | Cat# 13778100 |
| µ-Slide 8 Well | Ibidi | Cat# 80826 |
| Novex WedgeWell 10 to 20%, Tris-Glycine, 1.0 mm, Mini Protein Gel | Invitrogen | Cat# XP10205BOX |
| Novex 4X Bolt LDS Sample Buffer | invitrogen | Cat# B0007 |
| NuPAGE 3 to 8%, Tris-Acetate, 1.5 mm, Mini Protein Gel | Invitrogen | Cat# EA0378BOX |
| Power SYBR Green PCR Master Mix | Applied Biosystems | Cat# 4367659 |
| Protease Inhibitor Cocktail | Roche | Cat# 11697498001 |
| Protein A/G Agarose Beads | Thermo Scientific | Cat# 20421 |
| PVDF Transfer Membrane, 0.45 µm | Thermo Scientific | Cat# 88518 |
| RIPA Lysis Buffer, 10X | Millipore Sigma | Cat# 20-188 |

Software and Algorithms:

| Reagent or Resource | Source | Identifier |
|---|---|---|
| DESeq2 | Love et al., 2018.[8] | https://bioconductor.org/packages/release/bioc/html/DESeq2.html |
| FCS Express | De Novo Software | Research Version 7 |
| FIJI | NIH | https://fiji.sc/ |
| GSEA | Subramanian et al., 2005.[9] | Version 4.2.0 |
| Ingenuity Pathway Analysis (IPA) | Qiagen | N/A |
| MAGeCK | Li et al., 2014.[10] | https://sourceforge.net/p/mageck/wiki/Home/ |
| MetaXpress | Molecular Devices | N/A |
| NIS-Elements | Nikon | N/A |
| PRISM | GraphPad Software | Version 7 |
| Salmon | Patro et al., 2017.[11] | https://combine-lab.github.io/salmon/ |
| tximport | Soneson et al., 2015.[12] | https://bioconductor.org/packages/release/bioc/html/tximport.html |

#### Supplementary References

1 Haney, M. S. *et al.* Identification of phagocytosis regulators using magnetic genome-wide CRISPR screens. *Nature genetics* **50**, 1716-1727, doi:10.1038/s41588-018-0254-1 (2018).

2 Monaco, G. *et al.* RNA-Seq Signatures Normalized by mRNA Abundance Allow Absolute Deconvolution of Human Immune Cell Types. *Cell reports* **26**, 1627-1640.e1627, doi:10.1016/j.celrep.2019.01.041 (2019).

3 Dragich, J. M. *et al.* Autophagy linked FYVE (Alfy/WDFY3) is required for establishing neuronal connectivity in the mammalian brain. *eLife* **5**, doi:10.7554/eLife.14810 (2016).

4 Orosco, L. A. *et al.* Loss of Wdfy3 in mice alters cerebral cortical neurogenesis reflecting aspects of the autism pathology. *Nature communications* **5**, 4692, doi:10.1038/ncomms5692 (2014).

5 Filimonenko, M. *et al.* The selective macroautophagic degradation of aggregated proteins requires the PI3P-binding protein Alfy. *Molecular cell* **38**, 265-279, doi:10.1016/j.molcel.2010.04.007 (2010).

6 Fang, J. *et al.* Metformin alleviates human cellular aging by upregulating the endoplasmic reticulum glutathione peroxidase 7. *Aging Cell* **17**, e12765, doi:10.1111/acel.12765 (2018).

7 Fox, L. M. *et al.* Huntington's Disease Pathogenesis Is Modified In Vivo by Alfy/Wdfy3 and Selective Macroautophagy. *Neuron* **105**, 813-821.e816, doi:10.1016/j.neuron.2019.12.003 (2020).

8 Love, M. I., Soneson, C. & Patro, R. Swimming downstream: statistical analysis of differential transcript usage following Salmon quantification. *F1000Res* **7**, 952, doi:10.12688/f1000research.15398.3 (2018).

9 Subramanian, A. *et al.* Gene set enrichment analysis: a knowledge-based approach for interpreting genome-wide expression profiles. *Proceedings of the National Academy of Sciences of the United States of America* **102**, 15545-15550, doi:10.1073/pnas.0506580102 (2005).

10 Li, W. *et al.* MAGeCK enables robust identification of essential genes from genome-scale CRISPR/Cas9 knockout screens. *Genome biology* **15**, 554, doi:10.1186/s13059-014-0554-4 (2014).

11 Patro, R., Duggal, G., Love, M. I., Irizarry, R. A. & Kingsford, C. Salmon provides fast and bias-aware quantification of transcript expression. *Nature methods* **14**, 417-419, doi:10.1038/nmeth.4197 (2017).

12 Soneson, C., Love, M. I. & Robinson, M. D. Differential analyses for RNA-seq: transcript-level estimates improve gene-level inferences. *F1000Res* **4**, 1521, doi:10.12688/f1000research.7563.2 (2015).
